# KMT2D haploinsufficiency in Kabuki syndrome disrupts neuronal function through transcriptional and chromatin rewiring independent of H3K4-monomethylation

**DOI:** 10.1101/2021.04.22.440945

**Authors:** Michele Gabriele, Alessandro Vitriolo, Sara Cuvertino, Marlene F Pereira, Celeste Franconi, Pierre-Luc Germain, Daniele Capocefalo, Davide Castaldi, Erika Tenderini, Nicholas Burdon Bèchet, Catherine Millar, Tom Koemans, Nitin Sabherwal, Connie Stumpel, Monica Frega, Orazio Palumbo, Massimo Carella, Natascia Malerba, Gabriella Maria Squeo, Tjitske Kleefstra, Hans van Bokhoven, Susan J. Kimber, Siddharth Banka, Giuseppe Merla, Nadif Kasri Nael, Giuseppe Testa

**Author notes:** These authors share first authorship (Michele Gabriele, Alessandro Vitriolo, Sara Cuvertino).

## Abstract

Kabuki syndrome (KS) is a rare multisystem disorder, characterized by intellectual disability, growth delay, and distinctive craniofacial features. It is mostly caused by *de novo* mutations of *KMT2D*, which is responsible for histone H3lysine 4 mono-methylation (H3K4me1) that marks active and poised enhancers. We assessed the impact of KMT2D mutations on chromatin and transcriptional regulation in a cohort of multiple KS1 tissues, including primary patient samples and disease-relevant lineages, namely cortical neurons (iN), neural crest stem cells (NCSC), and mesenchymal cells (MC). In parallel, we generated an isogenic line derived from human embryonic stem cells (hESC) for the stepwise characterization of neural precursors and mature neurons. We found that transcriptional dysregulation was particularly pronounced in cortical neurons and widely affected synapse activity pathways. This was consistent with highly specific alterations of spontaneous network-bursts patterns evidenced by Micro-electrode-array (MEA)-based neural network. Profiling of H3K4me1 unveiled the almost complete uncoupling between this chromatin mark and the effects on transcription, which is instead reflected by defects in H3K27ac. Finally, we identified the direct targets of KMT2D in mature cortical neurons, uncovering TEAD2 as the main mediator of KMT2D haploinsufficiency. Our results uncover the multi-tissue architecture of KS1 dysregulation and define a unique electrical phenotype and its molecular underpinnings for the cortical neuronal lineage.

## INTRODUCTION

Kabuki syndrome (KS) is a rare multisystem neurodevelopmental disorder (NDD) mainly characterized by intellectual disability (ID) and craniofacial dysmorphism^1^. About 60-70% of cases are caused by loss-of-function variants in KMT2D^2^ (previously known as MLL2 and ALR in human, and Mll4 in mouse) that are referred to as KS type 1 (KS1, OMIM:147920), while about 10% of cases are caused by deleterious variants in KDM6A^3, 4^ (KS2, OMIM:300867). KMT2D is a histone 3 lysine 4 mono-methylase (H3K4me1)^5^, while KDM6A encodes for one of the two histone 3 lysine 27 trimethyl (H3K27me3) demethylase ^6, 7^. KMT2D and KMD6A associate together in the KMT2D/COMPASS (COMplex of Proteins ASSociated with Set1) complex, which also includes p300, responsible for H3K27 acetylation. The H3K4me1 modification marks transcriptional enhancers, whose activity is scored by the concurrent presence of H3K27ac^8, 9^. Transcriptional enhancers are functional units of the 3D genome that are responsible for the activation of gene expression in time and tissue-specific manner^8, 9^. The mechanism by which enhancers activate gene expression involves long-range chromatin interaction and histone modifications but the precise mechanisms through which enhancers make contact with promoters and induce gene expression are not fully elucidated^10^. The interconnection between deposition of histone modification and the 3D genome organization is indeed a topic of intense investigation, especially for understanding the mechanism of neurodevelopmental disorders, since loss-of-function of these genes often cause neurodevelopmental disorders^11, 12^. The importance of genes involved in enhancer regulation is especially underscored by the fact that loss-of-function of p300 causes Rubinstein-Taybi syndrome 2 (OMIM:613684), while loss-of-function of YY1, which is involved in mediating the looping between promoters and enhancers ^13, 14^, is responsible for Gabriele de-Vries syndrome^13, 15^ (OMIM:617557) and also features a defect in H3K27ac at enhancers^13^. Despite the crucial role of KMT2D/COMPASS complex in priming enhancers with H3K4me1, and making H3K27 ready for acetylation by removing existing H3K27me3, recent research showed that the catalytic activity of both KMT2D and KDM6A is not necessary for transcriptional control^16–19^. Indeed, research underscored that different mechanisms other than their catalytical activity are involved in their transcriptional regulation. For example, as structural scaffolds of the COMPASS complex and by recruiting the histone acetylase p30020.

Therefore, we hypothesized that KS1 etiology could be mainly traced back to an altered stoichiometry of the KMT2D/COMPASS complex. Consequently, to accurately study the molecular pathology of KS1 it is necessary to study the impact of heterozygous truncating mutations in human cellular models. At the state of the art, the most robust attempt of model KS1 was the employment of mouse model, in which *Kmt2d* has been engineered to carry a *βGeo* cassette instead of the catalytical domain^22^. And could still exert its catalytically independent function^16, 17^. Therefore, this model may fail to recapitulate the impact of truncating mutation, leaving open the necessity of studying the impact of altered stoichiometry of KMT2D/COMPASS complex. Moreover, given the lack of a human model to study the molecular etiology of KS1, here we report a disease modeling strategy to identify the molecular hubs downstream of KMT2D, to fully mimic its haploinsufficiency. We used patient-derived tissues, and iPSCs and engineered hESC from which we differentiated disease-relevant tissues mainly affected in KS1.

## RESULTS

### An integrated KS1 cohort of primary tissues and iPSC-derived lineages

Our multi-centric disease modeling cohort included the following disease-relevant cross-tissue set of samples from KS1 individuals who were clinically diagnosed by interdisciplinary teams and received a molecular diagnosis of protein-truncating mutations (Fig.1A, Fig.1B): i) dermal fibroblasts from 7 individuals and 5 sex-matched unaffected parents (Fig.1A); ii) peripheral blood mononuclear cells (PBMC) from 5 additional KS1 individuals and 3 unmatched controls; iii) induced pluripotent stem cells (iPSCs) from dermal fibroblasts of 5 KS1 individuals plus 3 sex-matched unaffected parents; iv) iPSC-derived neural crest stem cells and mesenchymal cells of 5 KS1 their sex-matched controls; v) iPSC-derived Ngn2 from 5 KS1 individuals plus 3 sex-matched unaffected parents. Furthermore, we generated an isogenic line representative of a *KMT2D* mutation by employing CRISPR/Cas9 to engineer a frameshift mutation in exon 48 of the hESC MAN7 line. (Fig.1C).

**Fig.1.**
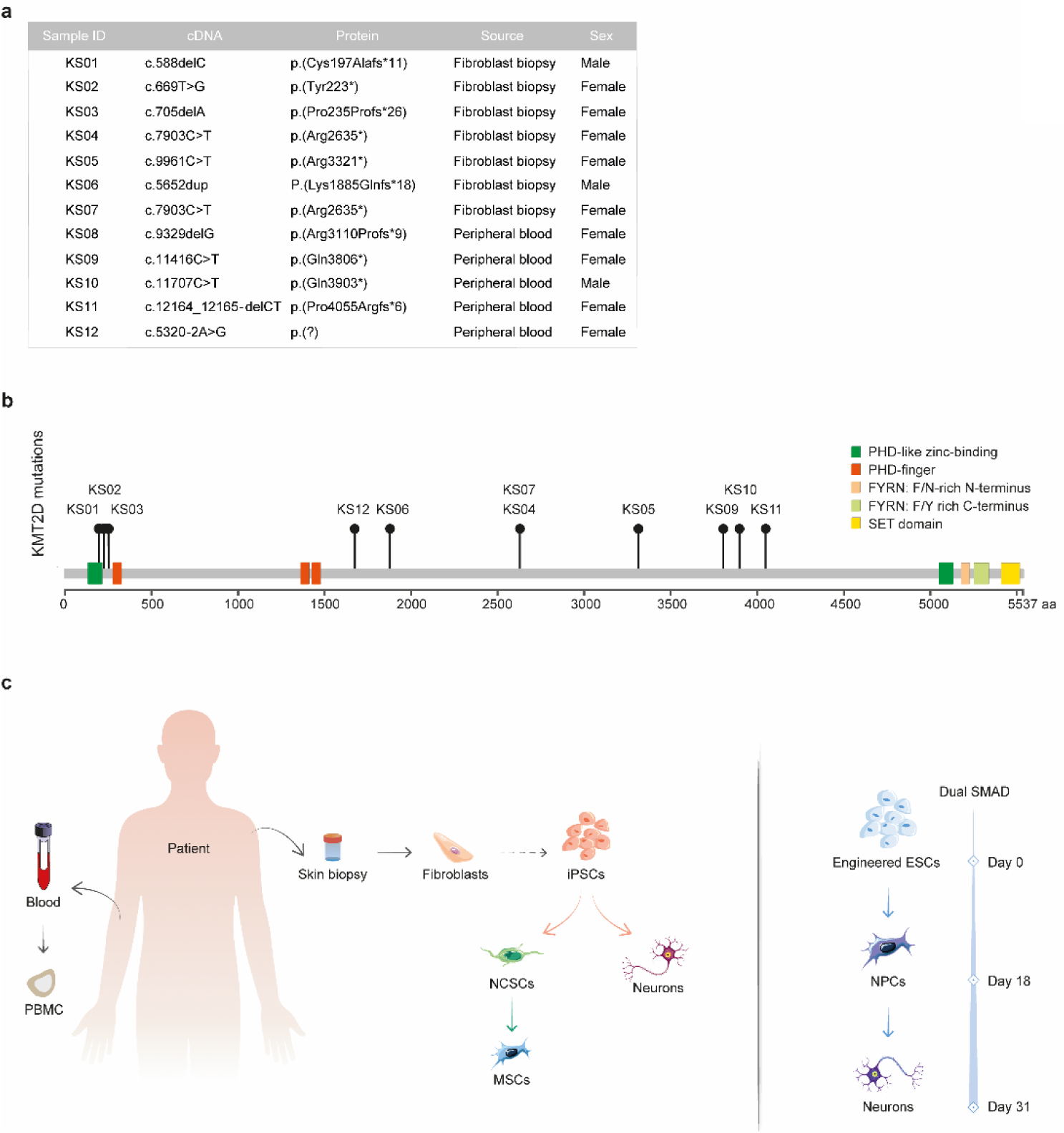
A) List of KMT2D mutations in KS1 individuals (<u>NM_003482</u>). B) Depiction of cohort mutations positions along *KMT2D*. C) Schematic drawing of our experimental design.

### Dermal fibroblasts and peripheral blood samples show disease-relevant dysregulations, minimally explained by vast changes in H3K4me1

Analysis of the transcriptome of dermal fibroblasts detected 185 differentially expressed genes (DEGs) (FC threshold=1.5, *FDR=0.05*). *KMT2D* levels were halved in most KS1 lines (*FC=1.64*, *FDR=0.08*, Fig.2A). Among altered transcripts we found a large set of genes (mostly upregulated) involved in neuron differentiation and related biological processes (BP) (Fig.2B), suggesting the presence of a *KMT2D*-specific signature in differentiated non-neural tissues, and allowing for the evaluation of patients’ fibroblasts as a good source for disease-relevant investigations. To identify those molecular hubs interposed between KMT2D and affected genes we performed a master regulator analysis. We found *EZH2* as the most enriched master regulator, accounting for 18.92% of DEGs, mostly upregulated (*p=1.36e^-^*^17^). Among enriched transcription factors (TF) we found *TEAD4*, which is upregulated in KS1 samples (*FC=1.51*, *p=0.002*) and it is found to be an upstream regulator for 58% of DEGs.

**Fig.2.**
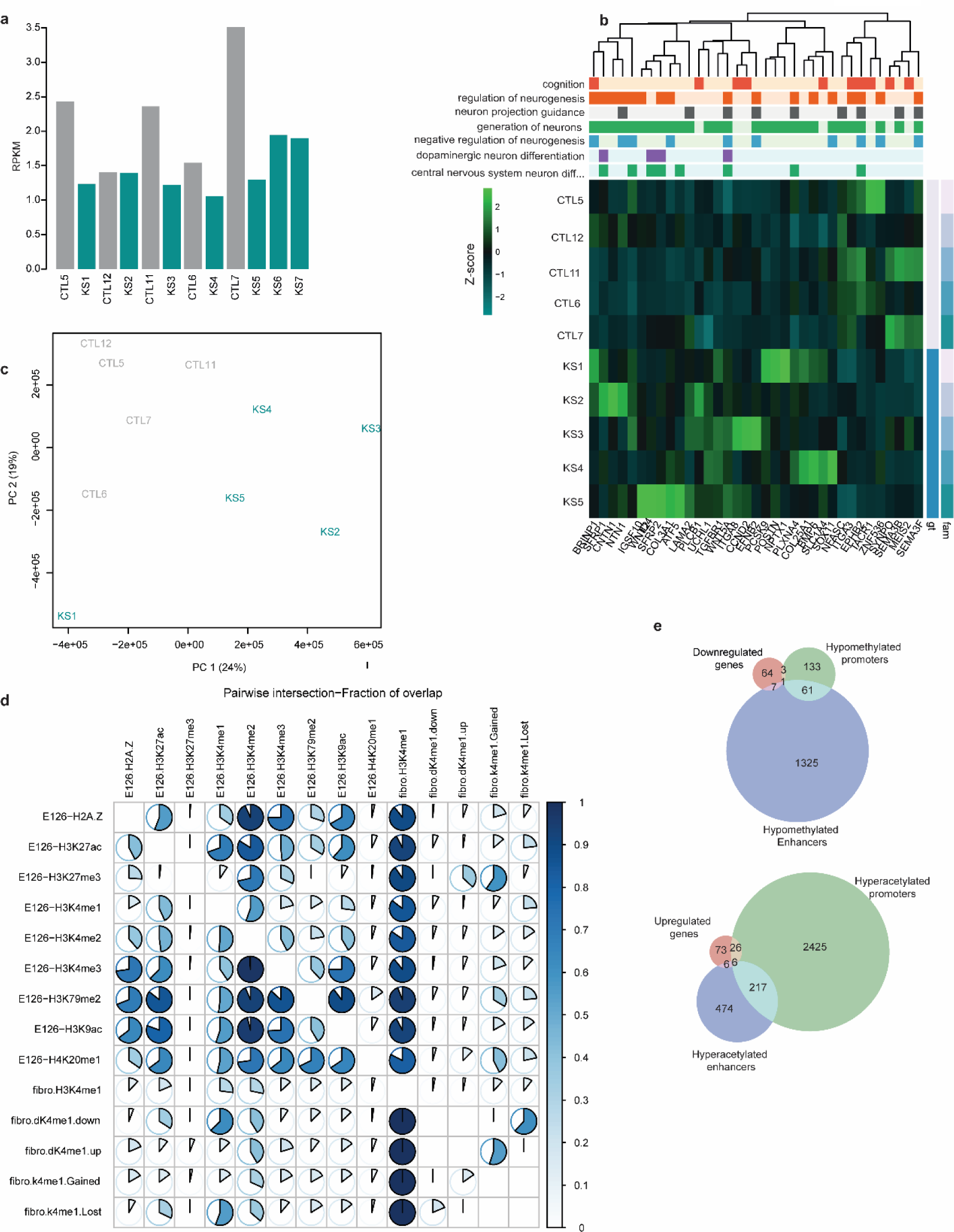
A) Barplot of *KMT2D* levels (RPKM); samples are coupled by family; controls are depicted in grey, KS1 in dark cyan. B) Heatmap of DEGs enriching GO categories (BP) associated with cognition and neurodevelopment; GO-categories are annotated on top of gene expression levels, reported as logTMM scaled by gene (z-score); family and genotype are annotated on the right. C) Principal component analysis of H3K4me1 read-counts (normalized on library size); points for control samples are labeled in black, KS1 in dark cyan. D) Pairwise intersection of diff-H3K4me1 regions with Roadmap Epigenomics; the fraction of overlap between couples of chromosomic location sets is reported as a circled heatmap; The last 5 sets identify regions that preferentially show H3K4me1 in KS1 (Gained), regions that do not show H3K4me1 in most KS1 (Lost), our reference of fibroblasts H3K4me1 peaks (fibro.H3K4me1), regions that show a stronger H3K4me1 signal in KS1 (fibro.dK4me1.up), regions that show a reduced H3K4me1 signal in KS (fibro.dK4me1.down). E126: adult dermal fibroblasts primary cells from Roadmap Epigenomics. E) Venn diagrams of DEGs and differentially methylated regions; upregulated and downregulated genes are divided, and respectively compared with hyper and hypomethylated genes; hypermethylated and hypomethylated enhancers refer to genes whose enhancers are quantitatively hyper or hypomethylated, the same logic was followed for hyper and hypomethylated promoters.

Next, to measure the impact of KMT2D haploinsufficiency in H3K4me1, we first probed its alteration in the 5 half-matched KS1 lines and relative controls. Western blot analysis did not reveal any difference in the bulk abundance of H3K4me1, H3H4me2, and H3K4me3 (Supplementary Fig.1A). Therefore, to identify local dysregulation, we performed H3K4me1 chromatin immunoprecipitation coupled with sequencing (ChIPseq). The principal component analysis (PCA) of H3K4me1 ChIPseq separates the samples through the first principal component (PC1) according to the genotype (Fig.2C). Overall, we detected a higher, albeit not significant, number of H3K4me1 peaks in KS1 individuals. Then, to identify *bona fide* gained and lost peaks in KS1 and to identify regions of the genome where the ChIPseq signal was significantly different between controls and KS1 we performed a qualitative and quantitative analysis, respectively.

Moreover, to compare our results with consolidated physiological datasets, we compared our H3K4me1 signal with Roadmap Epigenomics ^18^ (Fig.2D) and crossed H3K4me1 with the DEGs list to identify the amount of association between H3K4me1 and transcriptional dysregulation. The global distribution of H3K4me1 in our samples, as well as the differentially mono-methylated regions greatly overlapped with H3K4me1 and H3K4me2 identified by Roadmap Epigenomics (Fig.2D).

Regions with reduced H3K4me1 overlapped with H3K27ac regions, suggesting a loss of H3K4me1 at active regulatory regions. Regions gaining peaks or higher H3K4me1 signal largely overlapped H3K4me2 regions. Notably, only a fraction of downregulated DEGs (∼14%) was found to be affected by a change in H3K4me1 at regulatory regions (Fig.2E), while ∼34% of upregulated genes were affected (*p=2.18e^-05^*), and the largest portion of increased H3K4me1 signal was focused at promoters (Fig.2E).

In blood samples, we identified 193 DEGs (FC>1.5, FDR<0.05, Supplementary Fig.1B). DEGs were enriched in 7 biological process categories composed mainly of downregulated genes in KS cells (Supplementary Fig.1C). Remarkably, the “Cell fate commitment” category included DEGs whose function is also involved in central nervous system development (Supplementary Fig.1D).

### KS1 patient-derived induced pluripotent stem cells show only mild transcriptional and chromatin dysregulation

We reprogrammed eight fibroblast lines (5 from KS individuals and 3 from half-matched controls) to induced pluripotent stem cells (iPSC). Multiple clones for each iPSC were characterized for removal of reprogramming constructs and pluripotency marks (Supplementary Fig.2A and B). RNAseq studies on iPSCs based on 3-4 biological replicates reach the capacity to recapitulate differences between genotypes better than using multiple clones from fewer individuals ^22^. For this reason, we selected a single iPSC clone from each individual for the following experiments.

Differently from what was observed in fibroblasts, *KMT2D* expression in pluripotent lines did not mirror haploinsufficiency (Fig.3A). Differential expression analysis (DEA) found a largely reduced dysregulation compared to fibroblasts (63 DEGs *with FDR<=0.05*, *FC>=1.25*), showing no significant enrichment in any GO categories. The top TF enrichments were associated with SUZ12 and EZH2, consistent with the fibroblast data. These two proteins are the main subunits of Polycomb Repressive Complex 2 (PRC2).

**Fig.3.**
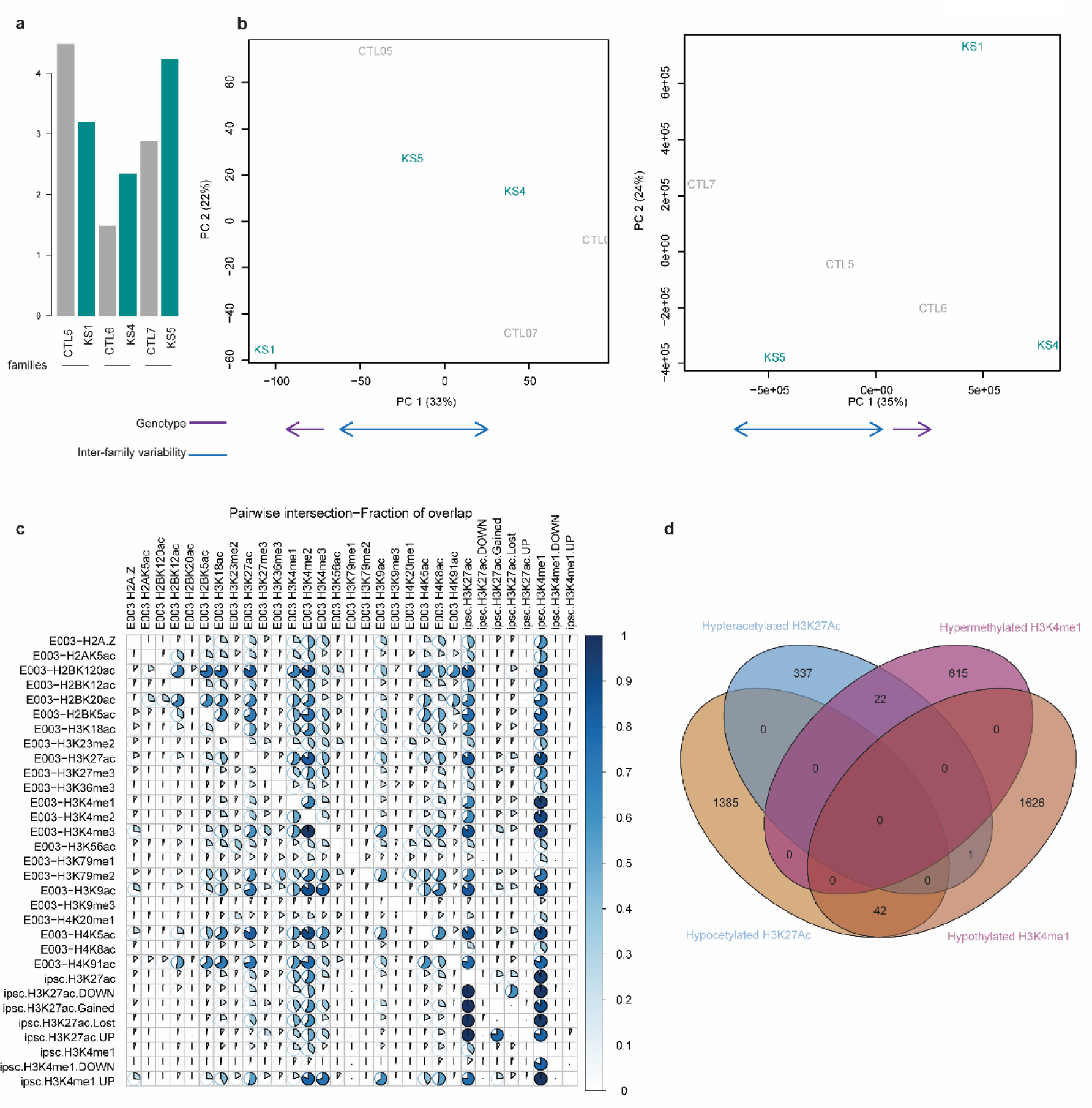
A) *KMT2D* is not differentially expressed in iPSCs. Barplot of *KMT2D* expression in our cohort (RPKM). CTL (grey) and KS1 (dark cyan) samples are coupled by family. B) PCA of H3K4me1 and H3K27ac (log-counts normalized on library size). A blue arrow depicts the observed inter-family variability, and a purple arrow depicts apparent intra-family genotype-dependent drifts. C) Pairwise intersection of iPSC reference (ipsc.H3K4me1 and ipsc.H3K27ac), differentially marked regions (.UP and .DOWN suffix), qualitative changes (.Gained and .Lost suffix) regions with Roadmap Epigenomics (E003-prefix); the fraction of overlap between couples of chromosomic location sets is reported with pie heatmaps. D) Venn diagram representing the intersection between iPSC differentially methylated (H3K4me1) and differentially acetylated (H3K27ac) regions; hyperacetylated regions are indicated in blue, hypoacetylated regions in yellow, hypomethylated in purple, hypermethylated in brown.

Given the enhancer-centric activity of KMT2D, we sought to assess the iPSC chromatin landscape by profiling H3K4me1 and H3K27ac using ChIPseq. Both marks showed a high inter-family variability on the PCA, with a mild but coherent intra-family shift, that could be associated with disease-specific differences (Fig.3B). Notably, compared with Roadmap Epigenomics data, our reference peaks completely included the equivalent marks (Fig.3D), meaning that we are sampling a large and comprehensive portion of both histone marks distribution. Moreover, H3K27ac peaks were completely included in our H3K4me1 locations, confirming that most H3K27ac is deposited in mono-methylated regions. Both quantitative and qualitative analyses revealed that most gains in H3K4me1 were found in regions that have enhancer activation marks, such as H3K18Ac and H3K27ac, and promoters such as H3K4me2 and H3K4me3 (Fig.3C). However, the extent of differentially H3K4me1 regions (15 regions passing *FDR=0.05*, 1667 passing *p=0.05*) in iPSCs was severely reduced compared to fibroblasts (∼10 thousand regions passing *FDR=0.05*), and no overlap can be found between regions differentially marked with H3K4me1 and dH3K27ac even upon lowering our significance threshold to *p=0.05* (Fig.3D). To further verify the limited dysregulation observed in iPSCs, we focused our attention on super-enhancers. They are clusters of distal regulatory regions that can be identified for their high and broad H3K27ac enrichment, usually associated with development regulation and cell-identity maintenance ^23^. Given the role of H3K4me1 and KMT2D in enhancer regulation, we hypothesized that super-enhancers could reveal a developmentally relevant dysregulation. Interestingly, we identified ∼10,000 super-enhancers. Among super-enhancers, we found a subgroup that was enriched in KS1 lines and showed a significant overlap with DEGs (Supplementary Fig.2D).

### The molecular disruptions of neurocristic axis caused by KS1 are evident only through differentiation

KS entails distinctive craniofacial traits ^24^. During early embryonic development neural crest stem cells (NCSC), migrate from the neural tube to populate the entire body, where they differentiate into mesenchymal cells (MCs) and further, to give rise to structures such as cartilages, bones, muscles, and secondary tissues, including those of the head ^25^. Consequently, we differentiated KS1 iPSC into NCSC and MCs.

RNAseq performed on both cell-types identified a dysregulation (27 genes in NCSC and 98 in MC passing *FDR=0.05*) mildly altered in NCSC (only 3 and 5 regions respectively passing *FDR=0.05*). Thus, we hypothesized that the subtle dysregulation observed at single multipotent states could instead be masking a developmental process that unwinds through differentiation. To answer this question, we designed a simple analytical strategy to apply to RNAseq data. We considered each cell type (iPSC, NCSC, and MC) as one of three consecutive stages of differentiation, dividing CTL and KS1 samples, and correcting by genetic background (∼individual + cell type).

This allowed us to identify the genes up or downregulated in most individuals, 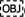consistent with that of CTL lines. These two groups were defined as “failed-up” and “failed-down” since they were not effectively up and downregulated in KS1 throughout simulated development (Fig. 4C and D).

**Fig.4.**
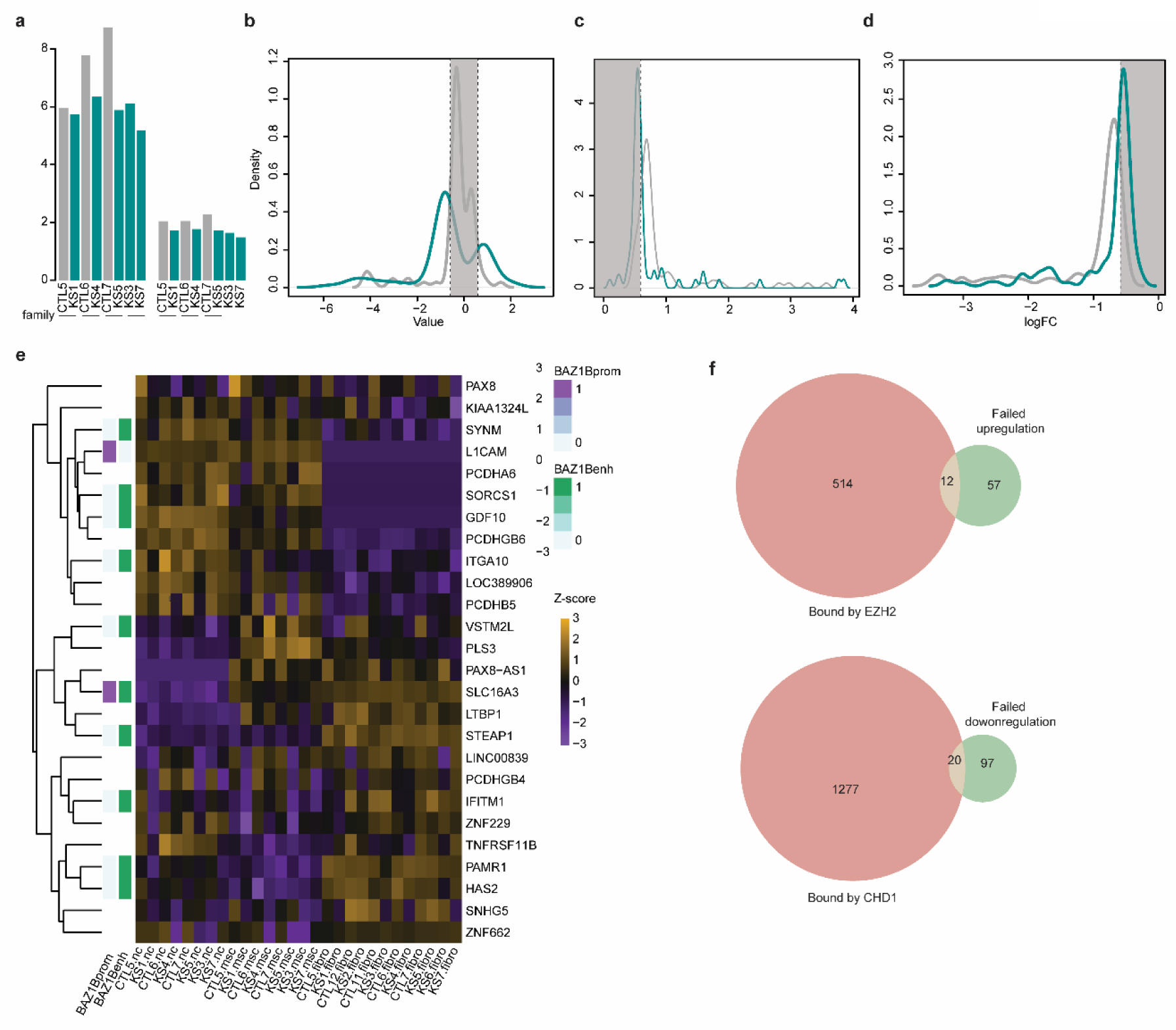
A) Barpot of *KMT2D* expression in both NCSC and MC. CTL (grey) and KS1 (dark cyan) samples are coupled by family. B) Kernel density distribution of logFC of genes differentially expressed along differentiation only in KS1 lines (darkcyan line and grey line respectively show KS1 and CTL logFC distribution for the same set of genes); a dashed vertical line reports FC=1.5. C) Kernel density distribution of logFC of genes upregulated along differentiation only in CTL lines («failed up» in the text; darkcyan line and grey line respectively show KS1 and CTL logFC distribution for the same set of genes); a dashed vertical line reports FC = 1.5 D) Kernel density distribution of logFC of genes downregulated along differentiation only in CTL lines («failed down» in the text; darkcyan line and grey line respectively show KS1 and CTL logFC distribution for the same set of genes); a dashed vertical line reports FC=1.5. E) Genes differentially expressed in at least two of the 3 neurocristic-axis cell-types; Heatmap of log-TMM scaled by row; Fibroblasts expression levels are also reported for comparison; row annotation reports known BAZ1B interaction with regulatory regions associated to the reported genes; BAZ1BProm refers to promoters bound by BAZ1B; BAZ1Benh refers to bound enhancers. F) Venn diagrams of the intersection between those genes that failed to increase the expression and those that failed to decrease the expression along the virtual development and known targets of the enriched master regulators.

Notably, we identified a significant enrichment for genes whose regulatory regions are bound by BAZ1B (Fig.4E), which is a master regulator of craniofacial development ^26^. Finally, we found two major enrichments for chromatin regulators in failed-up and failed-down genes, respectively corresponding to *EZH2* and *CHD1* (Fig.4F). Notably, *CHD1* mutations are known to cause Pilarowski-Bjornsson syndrome(OMIM: 617682), a neurodevelopmental disorder ^27^, exposing an interesting functional interaction with KMT2D.

### KS1 neurons have unique alterations in network activity

Intellectual disability is a hallmark of KS and one of its greatest burdens^28^. In order to elucidate its molecular underpinnings, we hypothesized that cortical neurons could reveal pathogenic vulnerabilities and differentiated iPSCs into excitatory cortical neurons through the transposon-based inducible *Ngn2* overexpression^29^ system that we recently validated^30^ (hereafter iNeurons). We performed morphological reconstruction on control and KS1 patient-derived iNeurons 23 days after *Ngn2* induction. Both control and KS1 neurons expressed upper layer markers (Fig.5A) and showed the expected neuronal morphology in terms of axonal and dendrite formation, as assessed by staining for AnkG and MAP2B (Fig.5B). To further characterize neuronal architecture, we performed low-density transfections to stain complete neuronal arborizations with dsRed and observed no morphological differences in neuronal architecture between control and KS1 for the number of primary dendrites, nodes, and dendritic endings (Supplementary Fig.3A-H).

**Fig.5.**
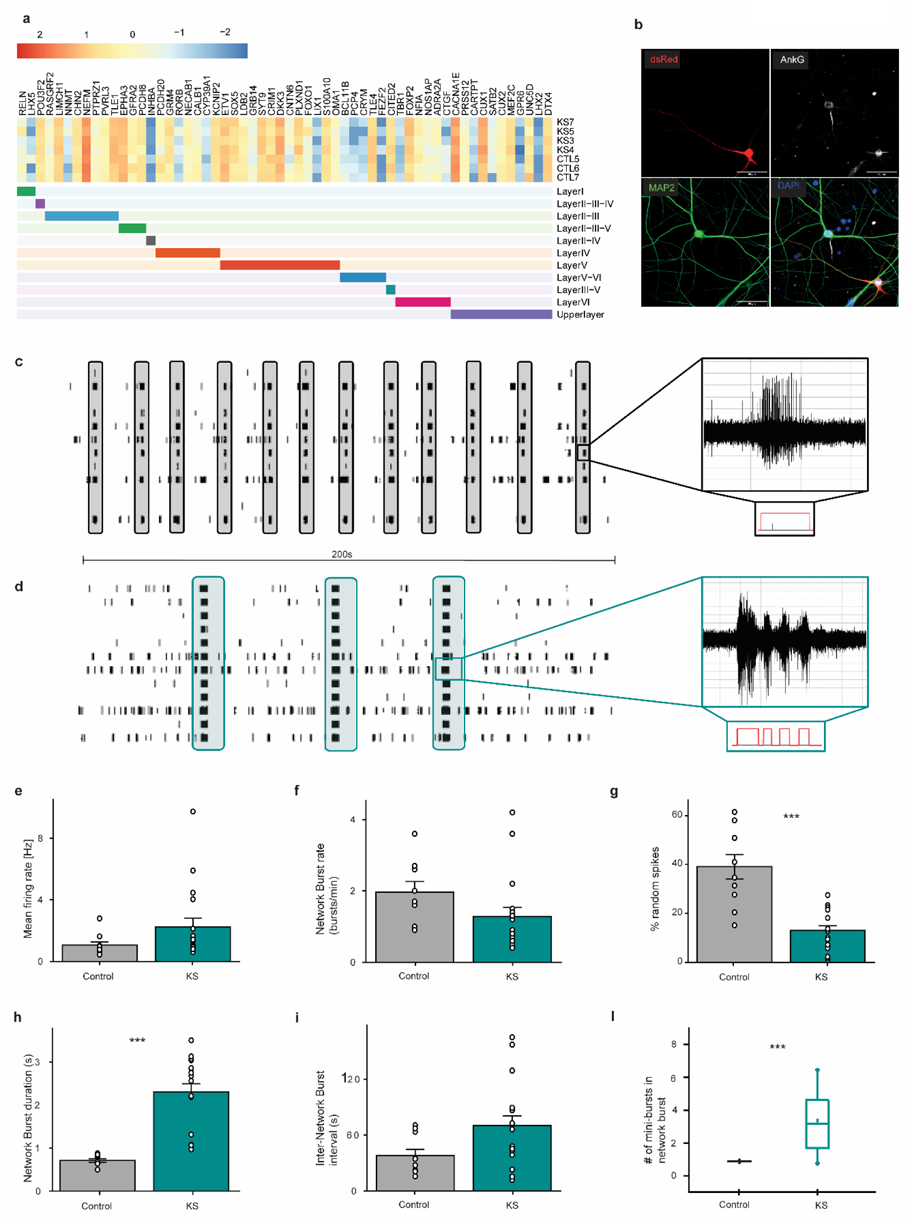
A) Characterization of neuronal identity. Expression profile of cortical layers markers. B) Neuronal architecture quality control. Representative image of immunostaining for AnkG (initial axon segment), MAP2 (dendrites), DAPI (nuclei), dsRed (entire neuronal structure). C) Representative raster plots showing 200 s recording of the spontaneous activity exhibited by control-derived neuronal networks during the fifth week *in vitro*. Network burst exhibited by the culture are highlighted in black. On the right panel, a representative single channel burst forming the network burst generated by control-derived neuronal network is shown (i.e. raw data of 3 s). D) Representative raster plots showing 200 s recording of the spontaneous activity exhibited by patient-derived neuronal networks during the fifth week *in vitro*. Network burst exhibited by the culture are highlighted in red. On the right panel, “mini-bursts” detected from a representative channel forming the network burst are shown (i.e. raw data of 3 s). E-L) Graphs respectively showing E) frequency of spikes (spike/s), F) frequency of the network burst, G) percentage of random spike, H) duration of the network burst, I) interval between consecutive network burst and L) number of “mini-burst” composing a network burst for neuronal networks derived from 4 control (n=10) and 4 KS1 patients (n=18) hiPSCs lines. Data represent means ± SEM. Statistics: two sample t-test, Kruskal-Wallis test, Bonferroni correction, p-values: *** = p<0.00025.

To identify potential alterations in identity and degree of maturation, we selected a curated panel of markers and analyzed their expression in KS1 compared to controls. We confirmed the cortical glutamatergic neuronal identity, with high expression of upper layer and mature neuronal markers (Supplementary Fig.4A). Next, we recorded the spontaneous electrophysiological network activity of iNeurons of either genotype grown on microelectrode arrays (MEAs) (Supplementary Fig.4B). Few days after plating, the neurons derived from healthy subjects formed functionally active neuronal networks, showing spontaneous events (i.e. spikes and bursts), as early as the second week of *in vitro* culture (Supplementary Fig.3J). Later in development (i.e. fifth week *in vitro*), the neuronal network showed a high level of spontaneous activity as well as regular network bursting pattern (i.e. synchronous events) involving almost all channels of the MEAs, indicative of a mature network (highlighted in grey in Fig.5C). KS1 iNeurons also established spiking activity during early network development as well as network bursts involving most of the channels of the MEAs later in development (Fig.5B). However, during the fifth *in vitro* week, KS1 patient-derived neuronal networks exhibited, at the population level, a global degree of activity that was slightly higher than controls (i.e. firing rate, Fig.5E, p=0.049). The level of synchronous activity exhibited by the KS1-derived neuronal network was comparable to control (Fig.5D, p=0.02). Interestingly, the pattern of activity exhibited by KS1 patient-derived neuronal networks was instead very different from controls. First, the percentage of random spikes (i.e. action potential events not organized within a burst ^31^) was significantly lower in KS1-derived neuronal compared to controls (Fig.5G, *p<0.00025*). Furthermore, the network bursts (i.e. synchronized trains of action potentials involving the whole neuronal network ^31^) appeared with longer durations compared to controls (Fig.5H, *p<0.00025*). This suggests that most of the activity in KS1 networks was organized in network bursts. Concentrating on these network bursts (see raw data highlighted in Fig.5C and D) we also found that the organization of the bursts within a network burst was sharply different. Specifically, while control network bursts were composed of single bursts appearing simultaneously in most of the channels, the KS1 network bursts were composed of “mini-bursts” (i.e. 4 “mini-bursts”, see raw data highlighted in red in Fig.5D), revealing an altered pattern of activity in KS1 excitatory neuronal networks. Together, these alterations outline a KS-specific electrical phenotype that is distinct from other NDD probed at the same resolution^32^.

### KS1 neurons have pervasive transcriptional dysregulation consistent with their electrophysiological phenotype

To identify the molecular underpinnings of the observed electrophysiological phenotype, we performed transcriptomic analysis on KS1 and control iNeurons at the same timepoint analyzed by MEA. By comparison to the other cell types, KS1 iNeurons showed a major transcriptional dysregulation yielding 1,181 high confidence DEG (FDR=0.05), including *KMT2D* itself that was precisely halved and *KDM6A* that was significantly upregulated (Supplementary Fig.5A). GO enrichment of DEGs reported molecular functions (MF) related to synaptic activity and ionic trafficking (Fig.6B), and biological processes (BP) involving memory and neuronal development (Fig.6C). Genes enriched both for “neuron” and “synaptic” categories, which might be responsible for the electrophysiological phenotype observed are depicted in Fig.6D. Among DEGs, we found a significant overlap with genes whose expression has been causally associated with several electrophysiological functions (EphysDEGs hereafter) in mouse neurons ^33^ (Supplementary Fig.5B).

**Fig.6.**
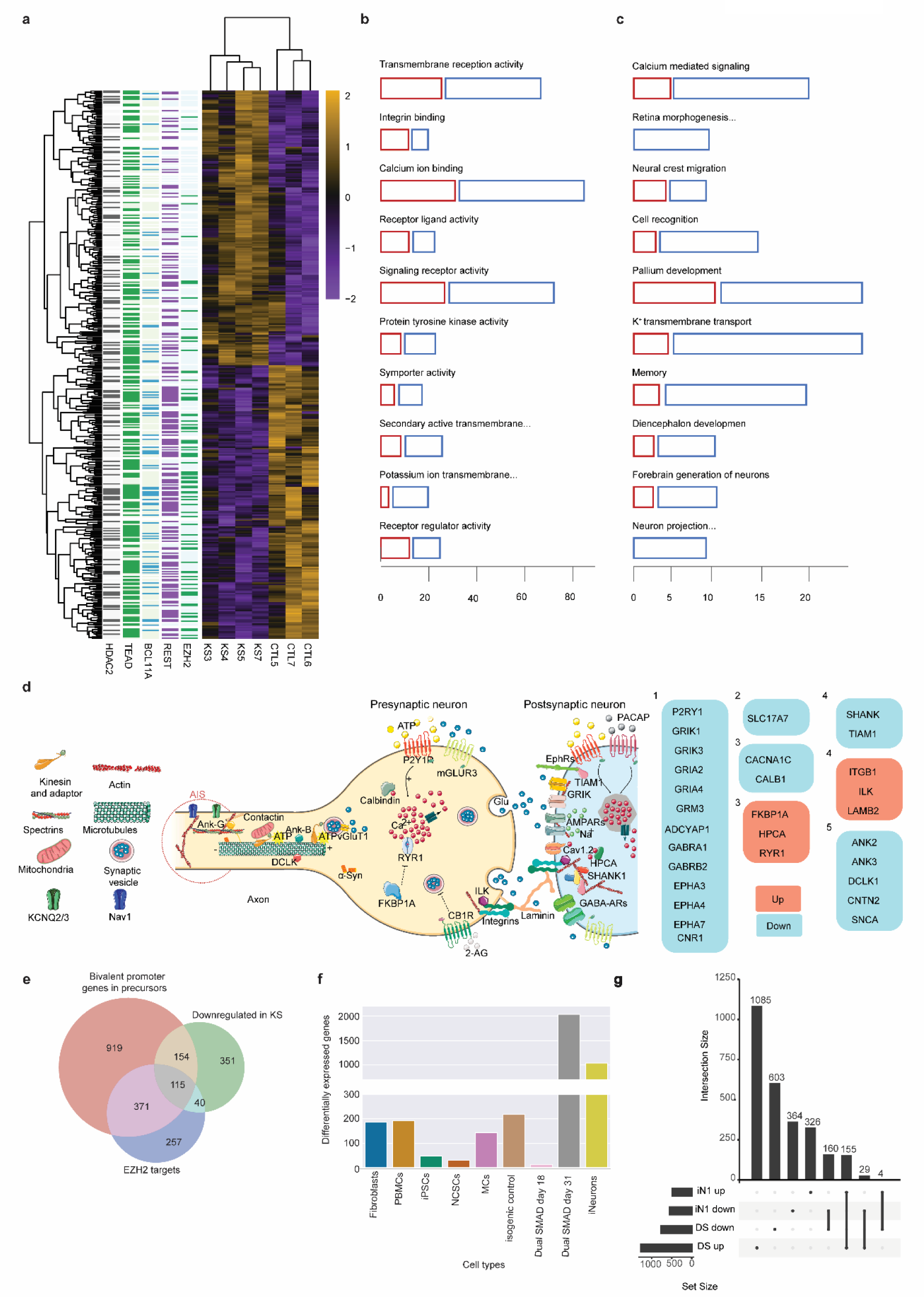
A) Heatmap of iNeurons (iN) differentially expressed genes (DEGs); logTMM scaled by row; targets of significant master regulators are annotated by row on the left. B) Molecular Function GO enrichments of iN DEGs; the horizontal axis represents the size (number of genes) of the enrichments, divided by sign of differential expression (blue for down and red for upregulated genes). C) Biological Process GO enrichments of iN DEGs; the horizontal axis represents the size (number of genes) of the enrichments, divided by sign of differential expression (blue for down and red for upregulated genes). D) Schematic depiction of iN DEGs enriching «neuron» and «synaptic» related GO categories. E) Venn diagram representing the overlap between genes bearing a bivalent promoter in NPC, genes downregulated in iN, and known targets of EZH2. F) Depiction of number of differentially expressed genes across analyzed tissues. In the isogenic setting and dual SMAD differentiation we identified the highest number of genes, followed by iNeurons. G) Representation of upregulated and downregulated genes both in DS and iN1, and corresponding intersections. H)

The most enriched master regulators we identified were *TEAD4*, *EZH2*, *REST*, *BCL11A*, and *HDAC2* (Fig.6A). *TEAD1*, *TEAD2*, and *TEAD3* were found upregulated, alongside *REST* and *VGLL4*, a partner of TEAD^34^. *TEAD4*, while upregulated in fibroblasts, was not expressed in iNeurons. Since TF target lists are mainly derived from motif enrichments and ChIPseq data, and all TEAD TF (*TEAD1*, *TEAD2*, *TEAD3,* and *TEAD4*) share the same DNA binding motif ^35^, we will generally refer to the TEAD family in our further analyses. Notably, EZH2 targets are preferentially enriched in downregulated genes, as observed also for the genes that failed to be up-regulated during differentiation down the neurocristic axis (failed-up genes.) We thus intersected downregulated genes in iNeurons with known EZH2 targets, and genes whose promoters are bivalent in neural epithelium ^36^, finding a highly significant intersection (p=1.20e-59, Fig.6E). Given the well-established role of EZH2 in neurogenesis and the bivalency characteristic of such DEGs promoters, this observation suggests that most downregulated genes in KS1 neurons are regulated by EZH2 and primarily involved in neuronal maturation.

In line with the benchmarking criteria we had empirically established for transcriptional endophenotypes scoring^22^, we pursued also an isogenic design to validate results obtained from patient-derived samples, generating a *KMT2D* mutant in the hESC MAN7 line using CRISPR/Cas9. We used this isogenic pair also to extend our results from *Ngn2* overexpression, which bypasses the early steps of neuronal differentiation, to the more developmental paradigm that generates cortical neurons by dual SMAD (DS) inhibition recapitulating major developmental milestones ^37^. The isogenic mutant line and the matched control were thus subjected to RNAseq in technical replicates prior to the onset of differentiation) and following, 18 and 31 days of dual SMAD differentiation and maturation, respectively representative of neuronal progenitors and cortical neurons. In this isogenic setting, KS1 transcriptional alterations were only noticeable at day 31, with the day 31 mutant line clustering at an intermediate distance between the control line at day 31 and all lines at day 18 (Supplementary Fig.5C). This separation in the principal component analysis suggests a delay in maturation, concordant with the observation of EZH2 downregulated targets in 5 weeks iNeurons and the mainly downregulated DEGs enriched GO developmental categories. Cortical neurons at day31, together with iNeurons, showed the largest dysregulation across tissues (Fig.6F). The intersection of DEGs in iNeurons and DS (FDR<0.05) was large and highly significant (195 genes, p=3.46e-81, Fig.6G), highlighting a core set of KS1 dysregulated pathways in terminally differentiated cortical neurons regardless of the protocol and/or developmental trajectory. Indeed, also in the isogenic line at day 31 of DS differentiation, we found the halving of KMT2D expression and the upregulation of TEAD genes and REST, corroborating the large overlap between the two neuronal models. *KDM6A* was instead not differentially expressed (Supplementary table 1), an observation that strongly down-plays the contribution of *KDM6A* upregulation to the transcriptional rewiring of KS1 iNeurons.

To further probe the altered maturation features of KS1 neurons, we performed a differential analysis of DS lines with an analytical strategy analogous to the one pursued for the neurocristic axis, i.e. using day 0, day 18, and day 31 as progressive stages, and identifying “failed-up” and “failed-down” genes along a virtual neuronal maturation path. Then, we intersected these genes with those physiologically upregulated during differentiation of cortical brain organoids(CBO), or preferentially expressed in neurons with respect to other cell types identified in the same CBOs ^38^. We thus identified a subset of approximately 250 genes whose expression changed coherently as differentiation of neurons progressed in two different human cellular models (CBO and DS). and found their regulation to be defective in *KMT2D* mutants, in both DS and iNeuron paradigms (Supplementary Fig.5D, Supplementary table 1). These genes, which constitute therefore the core subset disrupted in neuronal development in KS1, are enriched in GO biological processes such as regulation of synaptic transmission, molecular functions related to signaling and receptor activity, and cellular components such as post-synaptic membrane. Of particular note, within these categories, we found *SHANK1*, *GRIA2,* and *NEUROD1*, which have been respectively connected to autism spectrum disorder, neuronal electrophysiological activity, and neural maturity. Finally, among genes whose expression in CBO is directly correlated with *KMT2D* levels, we found two master regulators of KS1 iNeurons differentially expressed genes: *TEAD2* and *REST* (Supplementary table 1), corroborating the regulatory logic of KMTD2 observed in iNeurons.

### Alterations of H3K27ac largely explain transcriptional changes in KS1 neuronal models

Given the major transcriptional alterations identified in cortical neurons and the role of KMT2D in regulating enhancers, we profiled H3K4me1 and H3K27ac occupancy to define the impact of KMT2D haploinsufficiency on the enhancer regulome. Following the observations on MEA and RNAseq, we found that the vast majority of genes enriching “synaptic” GO categories were downregulated and hypo-acetylated at enhancers (Fig.7A).

**Fig.7.**
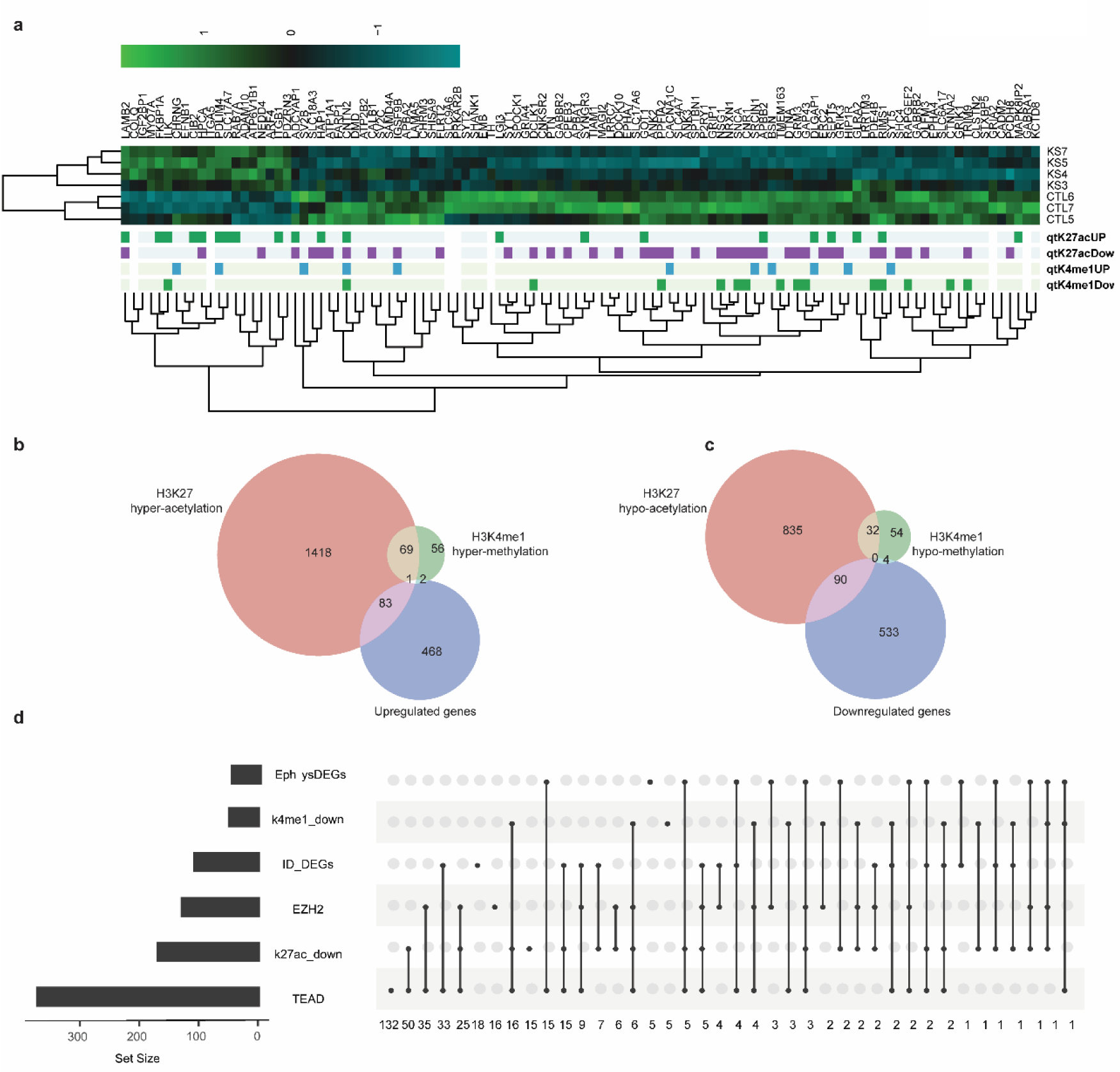
A) Heatmap of synaptic genes (logTMM scaled by column); differentially marked genes are annotated below; .UP and .DOWN suffix represent the direction of the dysregulation associated to enhancers of each gene. B) Venn diagrams showing the intersection between upregulated genes, and genes whose regulatory regions are hypermethylated or hyperacetylated (each list was filtered by FDR<0.05). C) Venn diagrams showing the intersection between downregulated genes, and genes whose regulatory regions are hypomethylated or hypoacetylated (each list was filtered by FDR<0.05). D) Simplified upset graph showing intersection of DEGs associated with ID (ID_DEGs), correlated with electrophysiological traits (EphysDEGs), regulated by EZH2 or TEAD, whose enhancers are hypomethylated (k4me1_down) or hypoacetylated (k27ac_down); number of genes accounting for each group is reported on the left; number of genes composing intersection of multiple groups are reported below each column.

Analysis of iNeuron chromatin marks revealed that changes in H3K27ac were larger and better able than H3K4me1 to explain changes in gene expression (Fig.7B, Supplementary Fig.6A). In particular, as expected from the H3K4me1 distribution prevalence at enhancers ^39^, almost none of the DEGs had changes in H3K4me1 at the promoter region (only *CHRNG, HHATL, PTGS1, and SNX33* with *FDR<0.05*). On the other hand, 17.79% of downregulated DEGs showed a significant (*FDR<0.05*) decrease in H3K27ac at promoters and 13,79% at enhancers. Strikingly, the number of hyper-acetylated regions (and associated genes) was larger than that of the hypo-acetylated ones (Fig.7B and C), conceptually in agreement with *KDM6A* upregulation. However, upregulated genes with increase H3K27ac at regulatory regions did not show any GO enrichment (Supplementary Table 1). This contrasts with downregulated genes with a concordant decrease in H3K27ac (Supplementary Table 1). Indeed, hypoacetylated downregulated genes largely showed enrichments for categories such as cell adhesion, neuronal differentiation, axon development, and neuronal migration (Supplementary Fig.6B).

These genes included 17 EphysDEGs (*not significant*) and 37 genes previously associated with ID (*p=1.52^e-^*^51^, Fig.7D). Notably, most downregulated genes losing H3K27ac at enhancers are known TEAD and EZH2 targets (77.19% and 30.99% respectively). Among ID-associated DEGs, 33.95% are differentially acetylated at enhancers (*p=2.67e^-29^*), and most of them are TEAD targets (70.27%).

Given the large dysregulation observed of H3K27ac, we further investigated for genomic locations that acquire or lose the status of “super-enhancers”, and identified DEGs associated with those regions (Supplementary Fig.6C). The overlap between genes associated with regions that lost the super-enhancer condition in KS1 and downregulated genes was larger than the opposite upregulated genes gaining super-enhancers in KS1. Notably, the downregulated subset included *KMT2D,* and the upregulated group included *VGLL4*, reinforcing the observation of a cross-talk between KMT2D and master regulators.

### Rescue of KMT2D and its downstream master regulators expression buffers KS1 transcriptional dysregulation in neurons

Given the defect in neuronal and synaptic maturation revealed by the transcriptional dysregulation observed in KS1 neurons, both in patient-derived samples and in the isogenic setting, we sought to determine to which extent this delayed phenotype could be overridden by accelerating neuronal differentiation through well-established media of defined composition (Neurobasal Plus, hereafter referred to as iN2). We thus repeated the transcriptional and chromatin analysis and found, as expected, that when compared to the iN1 condition, iN2 supplementation had a significant transcriptional impact on both controls and KS1 samples. KS1 lines were however much more affected than controls (Supplementary Fig.7A), with the number of genes differentially expressed in controls between the two conditions amounting to only one-fifth (Supplementary table 1) of the genes dysregulated in KS1 between the two conditions.

Notably, *KMT2D* was not downregulated in iN2. In addition, a significant subset of genes affected by iN2 in both CTLs and KS1, were differentially expressed in iN1 (between CTLs and KS1). This suggests that the majority of genes differentially expressed in iN1, as tightly regulated by *KMT2D*, are largely sensitive to environmental conditions and tightly linked to KMT2D genetic dosage (Supplementary Fig.7A, indicated as “∼gt CM” in Venn diagram), which ceased to be so in iN2 (*p< 0.01*).

Neuronal markers were similarly expressed across DS, iN1, and iN2 (Supplementary Fig.7B, Supplementary Table 1), with few markers specifically expressed in iN2, and some shared by iN2 and DS (Supplementary Fig.7C). GO enrichment for biological processes in iN2 DEGs referred to developmental processes not strictly linked to neuronal activity, while molecular function enrichment pointed towards chromatin and transcriptional regulation (Supplementary Table 1, GO tables), hinting at a dysregulation that does not pervade neuron-specific domains of regulation. When we compare the genes differentially expressed between KS1 and CTL lines across our neural models, we observed a strong overlap between DS and iN1, and a milder dysregulation in iN2 (Fig.8A). The overlap between neuronal markers expressed in the three different settings (iN1, iN2, and DS), is larger between DS and iN2. Concurrently, the overlap between DEGs is larger between DS and iN1. Thus - in a continuum of cell identities represented by iN1, iN2, and DS - being DS apparently more similar to iN2, and being most DEGs of iN1 rescued in iN2, we can consider iN2 media to be capable of rescuing most of DS dysregulations. Indeed, the expression of most master regulators identified in iN1 was rescued or reverted in iN2 (Supplementary Table 1). Moreover, a large portion of genes differentially expressed in DS and iN1 did not appear to be dysregulated in iN2, including most of the DEGs associated with ID (Fig.8B). When we compared the amplitude of dysregulation of DEGs (*FDR<0.25*) in iN1 and iN2, 56% of genes showed the same sign of dysregulation (*p=0.007*), with very few DEGs bearing opposite significant fold-changes (Fig.8C). Notably, iNeurons grown in iN2 showed a very mild dysregulation of genes transcriptionally correlated with neuronal electrophysiological properties (Supplementary Fig.7D), including the rescue of TEAD dependent dysregulation of EphysDEGs observed in iN1. In general, TEAD targets were less enriched by DEGs of iN2, and the main master regulators, whose targets were enriched, were *EZH2*, *CTBP2*, *SUZ12*, and *BCL11A* (*FDR<0.001* and enrichment>2).

**Fig.8.**
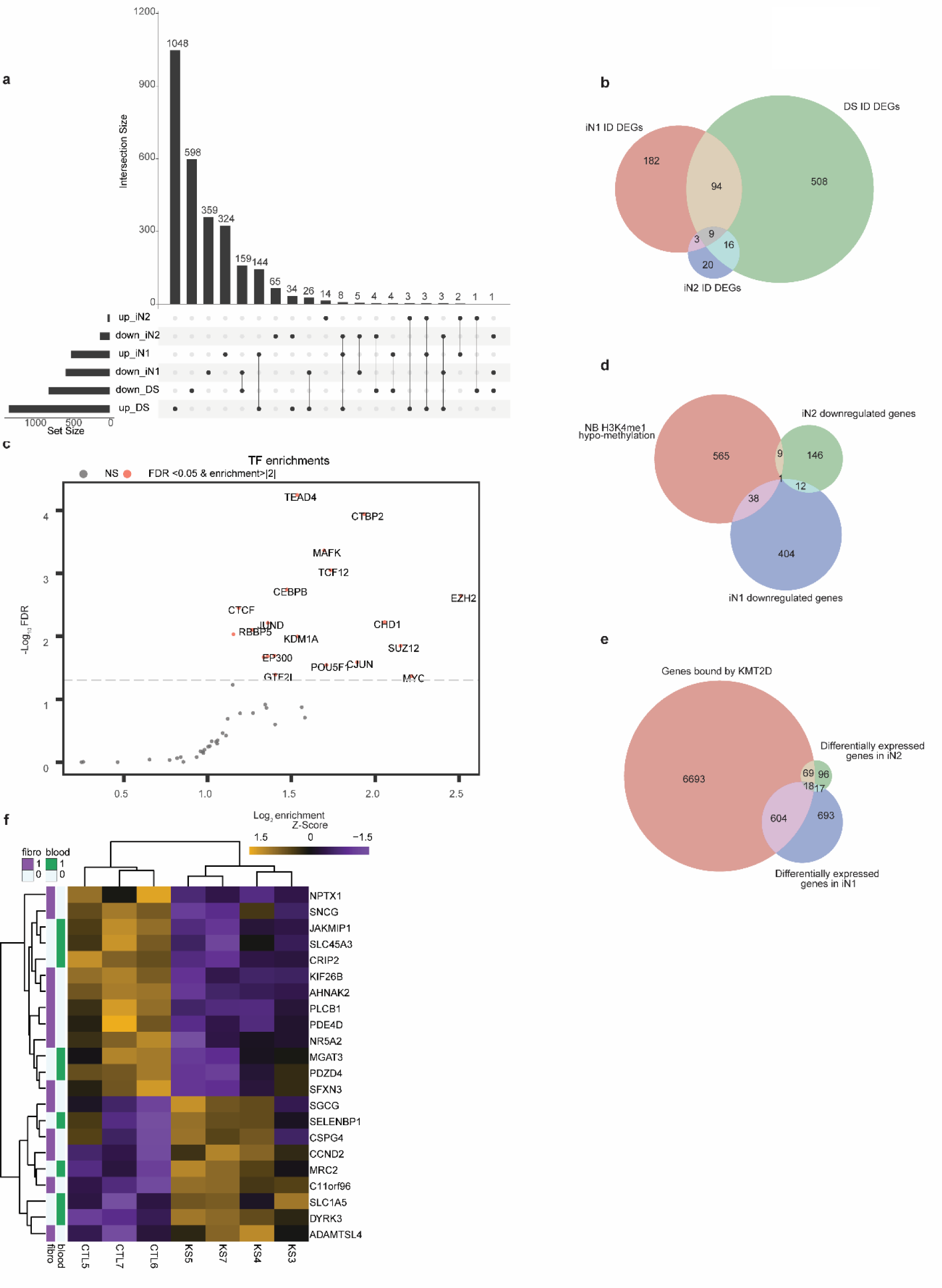
A) Upset diagram representing the number of genes differentially expressed in each neuronal differentiation culture. B) Venn diagrams representing the number of differentially expressed genes associated to ID in each neuronal differentiation culture. C) Transcription factors enrichments plot; TF whose targets are significantly enriched by DEGs bound by KMT2D at their enhancers in iN2. D) Venn diagram intersecting hypomethylated enhancers and genes downregulated in iN1 and iN2. E) Venn diagram intersecting genes bound by KMT2D at enhancers, with iN1 and iN2 DEGs. F) Heatmap of genes differentially expressed in iN1, which are also DEGs in fibroblasts or blood samples. Relative expression is expressed as z-scores of iN1 TMM levels. Rows are annotated on the left of the heatmap, to report the somatic tissues in which they are differentially expressed.

To identify the alterations of H3K4me1, H3K27ac, and H3K27me3 in iN2 we performed Cut&Run^40^, which allows an input of lower number of cells and improved signal-to-background ratio compared to classical ChIPseq. Notably, concordantly with the strong reduction of DEGs number, we did not observe a large significant change in H3K27ac, while we observed a higher dysregulation in H3K27me3 and an even larger dysregulation in H3K4me1 (Supplementary Fig.7D). H3K4me1 showed the largest dysregulation in fibroblasts and iN2 compared to H3K27ac, and the smallest in pluripotent and multipotent lineages, suggesting that H3K4me1 might be more important in more terminally differentiated cell types. Besides the numbers of H3K4me1 hyper- and hypomethylated regions were similar, the hypo-methylated regions largely overlapped regulatory regions, while the hyper-methylated regions could not be associated with genes. Moreover, as observed in the other tissues, the association between differentially H3K4me1 regions and DEGs was limited (Fig.8D).

To identify the direct targets of KMT2D we performed Cut&Tag^41^,, which is a more sensitive variant of Cut&Run. Most KMT2D bound regions overlapped both H3K4me1 and H3K27ac marks in both iN1 and iN2 (Supplementary Fig.7E). Most DEGs of both iN1 and iN2 were bound by KMT2D at enhancers (Fig.8E). Remarkably, we found KMT2D to bind its regulatory region and *KMT2A*, *KMT2B*, *KMT2C*, and *KMT2E* regulatory regions, as well (Supplementary Table 1).

Notably, we identified *EZH2* and *CTBP2* as master regulators associated with both groups. REST was strongly associated with iN1 DEGs, while *TEAD4* appears to have a higher enrichment in iN2 (Fig.8F and Supplementary Fig.8A). Interestingly, among genes bound by KMT2D and differentially expressed in both iN groups, we found *TEAD2* (with opposite FC in iN1 and iN2), which was also dysregulated in fibroblasts. Moreover, one-third (33,11%) of KMT2D targets were bound by TEAD (*p=0.037*) (Supplementary Table 1). This elects TEAD2, among other TEAD candidates, to be the main link between KMT2D and iN1 dysregulation reverted in iN2.

Since we observed a major transcriptional dysregulation mostly in terminally differentiated cell types (Fig.6F), we investigated whether the dysregulations found in peripheral blood, fibroblasts, and neurons converged on a critical subset of genes. We did not detect any overlap between differentially expressed genes in fibroblast and blood samples (Supplementary Fig.8B). On the other hand, both tissues shared dysregulated genes with iN1. Moreover, most of these genes dysregulated both in fibroblasts, blood, and iN1 were bound by KMT2D at enhancers (Supplementary Fig.8C), uncovering a subset of genes whose tissue-specific expression is vulnerable to *KMT2D* mutations (Fig.8F).

Finally, given the central role of both KMT2D and KDM6A in cancer^42, 43^, and the wide dysregulation observed in KS1 neurons, we investigated the transcriptional status of oncogenes ^44^ and tumor suppressors^45^ (Supplementary Fig.8D). Intriguingly, we found a significant overlap both for oncogenes (*p=0.005*), with 26 genes downregulated and 30 upregulated, and for tumor suppressors (*p=1.632e-06*), with 54 and 45 genes respectively down and upregulated. Moreover, both sets of genes showed enrichment in upregulated genes bound by KMT2D, which was significant with tumor suppressors (*p=0.04*).

## Discussion

One of the major opportunities of iPSC-based disease modeling for human genetics is the prospect of testing the impact of pathogenic mutations across multiple disease-relevant cell lineages, including those for which primary biopsies would be otherwise inaccessible, so as to define the multi-tissue molecular architecture that underlies multisystemic syndromes. Here we have actualized this potential for KS through a comparative design of primary and reprogrammed tissues, along with isogenic validation, to uncover the *KMTD2* dosage-dependent alterations across multiple lineages.

We found that mutant pluripotent and multipotent cell types showed only mild and very selective dysregulation. This is consistent with loss of function studies showing that KMT2D is necessary only at relatively late developmental stages (embryonic day E9.5, a crucial stage for brain and heart development ^46^), while also a murine model of KS1 showed the neural crest to be affected only after migration, at a terminal stage of differentiation ^47^. Amongst those we tested, cortical excitatory neurons were the most vulnerable cell type, with a major transcriptional dysregulation reflected in a prominent alteration of spontaneous synchronized electrical activity. Specifically, in KS1 neuronal networks, whose morphology and architecture were indistinguishable from controls, we identified prolonged and irregular network bursts when compared with both half-matched and independent controls. Accordingly, pervasive transcriptional dysregulation converged on electrical activity-critical pathways, as exemplified by *GRIA2* and *GRIA4*, respectively encoding AMPAR subunits 2 and 4, whose downregulation is consistent with the longer burst duration given the well-established properties of AMPA versus NMDA glutamate receptors-driven bursts (short and long, respectively) ^48^. Interestingly, the neuronal network phenotype included unique features (atypical burst shape consisting of many mini bursts) that distinguish KS1 from other neurodevelopmental disorders analyzed at equivalent functional resolution, as well as other properties (network burst rate and random spiking) that were similar to those observed in Kleefstra syndrome, a closely related neurodevelopmental disorder^32^, pointing to partially convergent mechanisms between the two disorders. As bona fide pathogenic endophenotype, such robust KS1-specific neuronal network properties are ideally suited for middle-to-high-throughput drug screenings. Given the major neuronal defect in H3K27ac deposition, these could be productively oriented towards histone deacetylase inhibitors, which have already proven valuable for a condition with a similar degree of intellectual disability (7q11.23 duplication syndrome, OMIM 609757)^30^.

At the chromatin level, transcriptional defects were aligned to the loss of H3K27ac, as we previously observed also in the context of Gabriele de-Vries syndrome, in which loss-of-function *YY1* mutations caused transcriptional alterations that matched loss of H3K27ac^13^. These results underscore, in the context of human KS tissues, the importance of the previously described coordinated action of the KMT2D-KDM6A-p300^20^ complex. Furthermore, we also confirm in the human setting previous observations^16, 49^ showing that KMT2D controls transcription independent of its catalytic activity. Indeed, we found the deposition of H3K4me1 mediated by KMT2D did not significantly overlay with transcriptional dysregulation, while the indirect impact on the histone mark for active enhancer H3K27ac was highly concordant with transcriptional deficits. By integrating transcriptomic and chromatin profiling alongside the genome-wide occupancy of KMT2D in the haploinsufficient setting, we can draw the following conclusions. First, we found KMT2D to bind the regulatory regions of the KMT2 family of methyltransferases commonly known as COMPASS complexes, as well as those of other key chromatin regulators and transcription factors, providing the molecular logic that explains the ramified gene network dysregulation caused by its decreased dosage. Indeed, KMT2D binds its own regulatory region and those of *KMT2A*, *KMT2B*, *KMT2C*, and *KMT2E,* highlighting a potential network of reciprocal regulation that becomes salient in light of the fact that mutations in all of them cause neurodevelopmental disorders with several shared features^50–52^. Second, our master regulator analysis revealed that *KMT2D* haploinsufficiency triggers precocious down-regulation of EZH2 targets. As EZH2, mutated in Weaver syndrome^53^ (OMIM: 277590), catalyzes H3K27me3, which is responsible for selective gene down-regulation to modulate neurogenesis at precise developmental stages 49, the observations that most down-regulated genes in all KS1 tissues we enriched in PRC2 targets hints at a dynamic competition between EZH2 and KMT2D for gene regulation during development. Third, one-third of KMT2D targets contained a TEAD binding site, pointing to an interplay between KMT2D/COMPASS complex and TEAD. This hypothesis is supported by the recent evidence of interaction between KMT2D and YAP/TAZ^54^, a known interactor of the TEAD family^55^. In particular, we found TEAD2 dysregulated both in fibroblasts and neurons, where it emerged as *KMT2D* direct target responsible for a vast portion of transcriptional dysregulation. Therefore, we suggest that the two proteins could act as complex, with TEAD driving DNA biding recognition of KMT2D. While the large size of KMT2D represents a challenge for interaction studies, our results ground the future investigation of a potential new regulatory axis in which TEAD2 is a partner of KMT2D/COMPASS and also regulated by it in a feedback loop. In such a scenario, the TEAD family could become a target for tackling KS1 manifestations as well as those cancer types in which KMT2D expression is altered. Indeed, given the importance of KMT2D in tumor biology ^56, 57^, we specifically focused on alterations in oncogenic pathways and found that in neurons several oncogenes and tumor suppressors were dysregulated by KMT2D haploinsufficiency, highlighting its central control of cancer-relevant cell identity across disease domains.

Finally, our characterization of the neurocristic axis allowed the identification of dosage-sensitive genes associated with craniofacial development^26^, which is critically and specifically affected in KS to the point of having originated its name. By placing the three multipotent lineages (iPSC, NCSC, and MC) on a virtual temporal axis we found that KTM2D haploinsufficiency resulted in subtle dysregulation that accumulated during development. Interestingly, however, while in animal models KMT2D was shown to be necessary for NCSCs formation and migration^58, 59^, we did not detect a major defect in this cell type, underscoring the value of iPSC-based modeling for defining the developmental stages of human-specific pathogenic significance.

In sum, our study elucidates across tissues then transcriptional and chromatin alterations of KS1, uncovering a major functional phenotype in neurons and defining the KMT2D dosage-dependent direct targets and intermediate effectors that underlie it. The combination, in this ample disease modeling cohort, of endophenotypes that are highly specific at the functional level and actionable at the molecular level, lays the foundation for rational drug screening studies. Finally, considering the KS1 enhancer dysregulation in light of the increasing relevance of long-range enhancer-promoter interactions^10^, future studies should focus on unveiling the impact of KMT2D loss-of-function on 3D genome organization, also in the context of the emerging potential of genome organization engineering for the treatment of human diseases^60^.

## MATERIAL AND METHODS

### Human samples

Participation in this study by patients and their relatives along with skin biopsy donations and informed consent procedures were approved by the ethics committee of Casa Sollievo della Sofferenza, San Giovanni Rotondo, Italy, protocol N.107/CE.

### Cell culture

Mycoplasma tests were performed routinely. Fibroblasts were cultured in RPMI 1640, 15% FBS, 1% L-Glutamine, 1% Penicillin-Streptomycin. Trypsin was used to passage fibroblasts. iPSCs were cultured with TeSR™-E8™ (Stem cell technologies) in feeder-free conditions on hES-qualified Matrigel (BD Biosciences) diluted 1:40. Three iPSCs clones were found mycoplasma positive and cleaned with Mycoplasma Removal Agent (Euroclone) prior differentiation. iPSCs were splitted with ReLSR (StemCell Technologies) or Accutase (Sigma) when single cells needed, supplementing the medium with rock inhibitor 5 μM Y-27632 (Sigma). NCSC were cultured following Menendez et al., 2013 ^61^. Cortical Neurons were induced and cultured as described elsewhere ^29^ with minor modifications. Cortical neuron maintenance was done with neurobasal medium fully complemented ^29^ and conditioned overnight on mouse astrocytes, or with Neurobasal Plus (Thermoscientific).

### Isogenic line generation by CRISPR/Cas9

#### Cell culture

hESC (MAN7_ctrl ^62^, MAN7_KMT2D-) were grown on Vitronectin (Life Technologies) using mTeSr1 medium (StemCELL Technologies). Cells were routinely passaged when 75-80% confluent using 0.1% EDTA-PBS without Ca^2+^ and Mg and were cultured at 37°C in a humidified 5% CO2 incubator.

### CRISPR-Cas9 gene editing

gRNAs (F: caccgCTATTGGTGAGAACAACGGG; R: aaacCCCGTTGTTCTCACCAATAGc) were designed to target exon 48 of human KMT2D using the Benchling software and cloned into pSpCas9(BB) vector. hESC cells (MAN7) were transfected with either KMT2D CRISPR-Cas9 vector or CRISPR-Cas9 empty vector together with GFP plasmid using P3 Primary Cell 4D-Nucleofector X kit (V4XP-3024, Lonza). Three days after transfection, GFP+ cells were sorted and plated on inactivated MEF cells. Colonies were then cut and passaged on feeder-free culture for amplification and subsequent analysis. Correct mutations were confirmed by Sanger sequencing.

### Dual SMAD differentiation protocol

hESCs were grown on Vitronectin (ThermoFisher Scientific). At the start of the protocol (adapted from ^63^) on Day 0 cells were plated on Matrigel (Corning) and cultured using mTESR1 medium (StemCell Technologies) supplemented by FGF2 (10ng/ml). On Day 1 neuronal differentiation was undertaken using neuronal induction medium supplemented by SMAD signaling inhibitors, Noggin and SD43154. On Day 10 cells were passaged and plated on a poly-l-ornithine and laminin (Sigma) substrate for neuronal maintenance and maturation. On Day 12 small-elongated cells generate rosette structures resembling early neuronal tubes and were propagated in culture using FGF2 (20ng/ml) for about 3 days. On Day 15-18 cells were passaged using Dispase (StemCell Technologies) and plated on a poly-l-ornithine and laminin substrate for neuronal maturation and cultured up to Day 31. Cells used for RNAseq were sorted for CD44^-^ CD24^+^ and CD184^+^ for the progenitor stage at Day 18 and for CD44^-^ CD24^+^ and CD184^-^(BD Bioscience) for neurons at Day 31.

### Reprogramming into iPSCs

Samples KS01, KS04, KS05, CTL_5, CTL_6, and CTL_7 were reprogrammed with self-replicating mRNA Simplicon™ RNA Reprogramming Technology (Millipore) according to the manufacturer’s protocol. Clones were tested by RT-qPCR for mRNA reprogramming construct removal ^64^ after B18R withdrawal. Samples KAB_3 and KAB_7 were reprogrammed using Sendai virus (CytoTune™-iPS 2.0 Sendai Reprogramming Kit, ThermoFisher Scientific) according to the manufacturer’s protocol. Sendai virus depletion was checked by RT-qPCR according to manufacturer’s instructions.

### SNP array

All fibroblast samples and iPSCs clones were subjected to SNP array for detection of genomic instability (Supplementary Table 2). High-resolution SNP-array analysis was carried out by using the CytoScan HD array (Thermo Fisher Scientific, Waltham, MA, USA) as previously described ^65^. Data analysis was performed using the Chromosome Analysis Suite Software version 4.1 (Thermo Fisher Scientific, Waltham, MA, USA) following a standardized pipeline described in literature ^66^. The clinical significance of each rearrangement detected has been assessed following the American College of Medical Genetics (ACMG) guidelines. Base pair positions, information about genomic regions and genes affected by copy number variations (CNVs), and known associated disease have been derived from the University of California Santa Cruz (UCSC) Genome Browser, build GRCh37 (hg19).

#### iPSCs differentiation into NCSC, MCs, and cortical neurons

NCSC differentiation was performed by treating iPSCs with small molecules GSK inhibitor and SB431542 as already described and further differentiated into MCs ^61^. NCSC differentiation was checked by fluorescence-activated cell sorting (FACS) using antibodies against NK-1 and NGFR, as previously described ^67^. MCs identity was verified by FACS analysis: CD44 (BD pharmingen, cat. No. 560531) and CD73 (BD pharmingen, cat. No. 344015) as previously described ^67^.

#### iPSCs differentiation into cortical neurons – iNeuron protocol

Cortical excitatory neurons were obtained by *Ngn2* overexpression in iPSCs. Pluripotent cells were engineered using an all-in-one ePiggyBac (ePB) transposon. We cloned the sequence of *Ngn2 –* P2A – EGFP – T2A – PuroR ^29^ sequence into an ePB backbone containing a blasticidin resistance cassette and doxycycline responsive element under the control of hUbC promoter ^30^. iPSCs were electroporated with the Neon system (ThermoFisher) with the following parameters: 1200V, 30 ms, 1 pulse using 0,5 µg of a helper plasmid expressing a transposase and 4,5 µg of donor plasmid with a transposable element. Electroporated cells were plated in presence of rock inhibitor 5 μM Y-27632 (Sigma). To isolate stably engineered lines, from the day following electroporation, cells were selected by the administration of blasticidin 5µg/mL. Neuronal differentiation was achieved by *Ngn2* overexpression, driven by doxycycline administration. Differentiation was performed as already described with minor modifications ^32^. Neuronal maturation was protracted up to five weeks. Stainings on neurons were performed using coverslips nitric acid-treated and coated overnight with poly-L-Lysine at 37°C, following human laminin coating (LN511-02 Biolamina), 3 hours at 4°C.

#### Immuno-fluorescence stainings

Prior RNAseq, Pluripotency of reprogrammed samples was addressed by TRA-1-60 live-cell imaging DyLight 488 conjugated (Stemgent, 09-0068), and NANOG (Everest Biotech, EB06860) in fixed cells. Stainings on fixed cells were performed as follows. Samples were washed with PBS and treated for 10 minutes with 4% Paraformaldehyde/4% sucrose, washed twice with PBS, and permeabilized with 0.2% Triton X-100 in PBS. One hour of blocking was performed in serum matched with secondary antibody specie. Following PBS washing, samples were incubated 2 hours at room temperature or overnight at 4°C with the primary antibody diluted in blocking buffer. Primary antibodies were removed by washing three times with PBS, and secondary antibodies were incubated for 1 hour at room temperature. Secondary antibodies were washed out with PBS. Samples were then treated for five minutes with DAPI, washed with water, and mounted on cover slides with Moviol mounting medium. Neuronal stainings were performed using, MAP2B (Abcam, Ab32454) and Ankyrin-G (N106/36, Millipore, MABN466).

#### Protein extraction and immunoblotting

RIPA buffer (150 mM NaCl, 1.0% NP-40, 0.5% sodium deoxycholate, Protease inhibitor cocktail (Sigma), 0.1% SDS and 50 mM Tris, pH 8.0) was used to lysate cells. Lysates were sonicated using the Bioruptor Sonication System (UCD200) for three cycles of 30 seconds at high power with 30 seconds pauses and centrifuged at 13,000g for 15 min. Bradford protein assay (Bio-Rad) was employed to quantify proteins. SDS-PAGE for histone modifications was performed loading 20–40 μg of protein extracts in home-made 10% polyacrylamide gels. Protein transfer was performed for 1 hour at 120V onto nitrocellulose membranes, which were blocked in 5% milk-TBS-T (50 mM Tris, pH 7.5, 150 mM NaCl, and 0.2% Tween-20). Antibodies were diluted in blocking buffer. Protein signal was quantified using the Odyssey Infrared Imaging System (LI-COR Biosciences). Densitometry was performed using ImageJ^68^. Secondary antibodies were α-rabbit IRdye680LT and α-mouse IRdye800LT (LI-COR Biosciences).

### RNAseq analysis

Expression levels for each RNAseq experiment were measured by quantifying reads with Salmon 1.0 quasi-mapping approach on a hg38 human transcriptome as done previously ^26^. Differential expression analysis (DEA) was performed with edgeR using the edg1 function described in (BAZ1B ref) when dealing with less than 3 individuals per genotype, edg2 was used in the other conditions. In the iPSC case, we aggregated reads, by sum, by individual, to obtain the 3 vs 3 setting described in the results section.

Neurocristic Axis DEA. To identify genes which activation or downregulation along development failed we separated iPSC, NSCS, and MC data by genotype, creating two separate RNAseq datasets. Then we performed a DEA using edg2, with ∼individual+celltype as model matrix. This allowed measuring gene expression fold-changes going from iPSC to MC. First, we identified genes whose expression changed along NCA only in KS1 lines (showing *FC<=1.5* and *or FDR>=0.25* in CTL lines; Fig.4B). Failed up-regulation was identified for genes whose FC was significantly positive (*FDR<=0.05*, and *FC>=1.5*) along the Neurocristic Axis (NCA) in CTL lines, while having a positive FC and *FDR>=0.25*, neutral or negative FC in KS1 lines (Fig.4C). Failed downregulation was identified for genes who had opposite behavior (significant negative FC in CTL, n.s. positive, neutral, or positive FC in KS1; Fig4D).

Cerebral Brain Organoids single-cell data analysis: Raw data was aligned with CellRanger v3.0 and only barcodes of cells that were assigned to one of the 15 cell-types identified in the original paper were retained. First, we aggregated counts by individual, excluding the “unknown” and “choroid” clusters. Taking into account gene-signatures and cluster coordinates into the UMAP of the original paper, we built 3 individual-specific mini-bulk datasets by summing read-counts of i) cluster 0,1, and 5 (Neurons and Early Neurons), ii) cluster 2,4,7,12 (Intermediate Progenitors and Outer Radial Glia) and iii) cluster 8,13,14 (Radial Glia). Thus, we obtained a dataset with 4 biological replicates of 3 virtual developmental stages. DEA analysis was performed using edg2, testing different stages as factors and correcting by individual (∼individual+stage).

### ChIPseq epigenomic analysis

ChIPseq was performed as previously described ^13^. Briefly, Chromatin cross-linking was performed using 1% formaldehyde in PBS and quenched adding glycine to the final concentration of 125mM. Cells were collected with ChIP SDS buffer (0.5% SDS, 5 mM EDTA, 100mM NaCl and 50 mM Tris-HCl at pH 8.1), centrifuged 400g for 30 min and resuspended in IP buffer (0.5% SDS, 5 mM EDTA, 100mM NaCl and 50 mM Tris-HCl at pH 8.6, 1,5% Triton X-100). Chromatin was sonicated with Branson digital sonifier (Emerson Industrial Automation) to obtain bulk DNA fragments of 300 bp. Chromatin for ChIP was quantified using Bradford protein assay (Bio-Rad). Immunoprecipitations were performed using 100 µg of chromatin. Antibodies used were: Ab8580 (H3K4me3), Ab8895 (H3K4me1), Ab4729 (H3K27ac), 9733B (H3K27me3). Libraries were prepared as already described ^69^ with adaptations for the automated system Biomek FX.

We performed ChIPseq reads alignment on the human hg38 genome using Bowtie 1.0 (-v 2 - m 1). Peak calling was performed with MACS2, using broad parameters (--broad –broad-cutoff 0.1 -q 0.1). To perform ChIPseq quantitative analysis, including PCA, we selected the peaks of each mark identified in at least two samples (independently of their genotype) and measure read-counts per peak in each sample with DeepTools multiBamSummary. On the resulting count data, we performed differential analysis with edgeR, normalizing on library size (number of reads mapped on the entire genome per samples), as previously described (ref BAZ1B). Differentially marked locations passing p=0.05 and FC=1.5 thresholds were considered significant in all tissues but fibroblasts and mature neurons, where we could found the largest differences by genotype, and we applied an FDR=0.05 significance threshold. To perform qualitative analyses we grouped i) peaks found in all samples of a certain genotype (e.g. KS1), that were present in at most 1 sample of the other genotype with ii) peaks found in at least n-1 samples of a certain genotype and not present in any of the other genotype samples. Both groupings were performed with BedTools (version 2.26) multiIntersect function.

### Roadmap Epigenomics comparison

We selected representative tissues and gathered all peaks passing FDR 0.1, for all marks, excluding those showing less than 500 peaks in a certain cell type. Specifically, we used E126 to compare with dermal fibroblasts, E003 for iPSC, E007 for NPC, E009 for neuronal progenitors, E071 for cortical neurons.

### Identification of regulatory regions

Promoters were defined as regions laying at −500/+250bp distance from transcription start sites (TSS). Enhancers were defined as regions showing H3K4me1 and H3K27ac signals, missing H3K27me3 and H3K4me3 signals devoid of TSS, as previously published ^26^ when in-house data or Roadmap Epigenomics ^70^ data were available, or downloaded from 4DGenome ^71^ and Psychencode ^72^, as indicated in the text.

### Identification of super-enhancers

Super-enhancers were identified following the ROSE code. We identified KS1 and CTL specific super-enhancers by selecting those found in all samples of each genotype, and then identified overlaps and enrichments by intersecting them with a set of super-enhancers found in at least two samples (independently of their genotype). Genes were associated with super-enhancers if their promoter was included or their enhancers were intersecting SE locations_73,74._

iNeurons super-enhancers definition: First, we relied on the ROSE pipeline as per the other tissues, to label H3K27ac regions in each sample as classic “enhancer” and “super-enhancer”. CTL-specific super-enhancers (CTL SuperEnh) were initially defined as regions called “super-enhancer” in at least 2 CTL samples (n-1), while KS1-specific ones (KS1 SuperEnh) were similarly defined as regions called “super-enhancer” in at least 3 KS1 (n-1) samples. Then we associated genes with CTL-and KS1-superenhancers, and removed genes spuriously associated with both groups.

### CUT&RUN and CUT&TAG

CUT&RUN and CUT&TAG were performed as previously described ^40, 41^ with minor modifications. In particular, CUT&TAG in neuronal samples was performed by dissociating neuronal culture with Accutase for 10 minutes followed by vigorous mechanical dissociation. Clumps were removed with a cell strainer and samples were centrifuged at 100g to remove debris, and at 200g to isolate the pellet enriched for nuclei. KMT2D antibody was donated by Kai Ge. CUT&RUN reads were aligned with Bowtie2 v2.2.5 as in ^40, 41^. Peak calling has been performed with Seacr 1.1 using relaxed threshold against IgG.

### Differential expression analyses

Differential expression analyses (DEA) were performed with EdgeR ^75^ with the parameter “estimateGLMRobustDisp”, taking into account genotypes, sex, family (when possible), and batches when present. Gene ontology analyses were performed by Goseq ^76^, Ingenuity pathway analysis (IPA, Qiagen), and the online tool webgestalt^1^, which permits to load a custom universe of genes as background (taking in account the expressed genes for each specific cell type) selecting Over-Representation Analysis.

#### Microelectrode arrays recordings and neuronal network analysis Neuronal differentiation

Eight iPSCs (4 lines reprogrammed from somatic cells of healthy subjects and 4 lines from KS1 patients) were directly derived into upper-layer excitatory cortical neurons by overexpressing the neuronal determinant neurogenin 2 (*Ngn2*) upon doxycycline treatment. The derived neurons were plated onto MEAs or glass coverslips pre-coated with adhesion-promoting factors (50 µg/mL Poly-L-Ornithine and 20 µg/mL Laminin) at a final density of 100 and 600 cells/mm2 respectively. Two days after plating, glia cells were also added to the culture at the same density. Differentiation was completed as described above.

### Morphology analysis

Cells plated on coverslips were transfected with plasmids housing Discosoma species red (dsRED) fluorescent protein 7 days after plating. After 23 days *in vitro*, cells plated on cover slips were fixed and mounted for imaging. Neurons were imaged using an Axio Imager Z1with 568nm laser light and a Axiocam 506 mono. Neurons were digitally reconstructed using Neurolucida 360 software (MBF–Bioscience, Williston, ND, USA).

### Micro-electrode array recordings

Recordings of the spontaneous activity of hiPSCs-derived neuronal networks were performed during the fifth week *in vitro*. All recordings were performed using the 24-well MEA system (Multichannel Systems, MCS GmbH, Reutlingen, Germany). MEAs devices are composed by 24 independent wells with embedded micro-electrodes (i.e. 12 electrodes/well, 80 µm in diameter and spaced 300 µm apart). Spontaneous electrophysiological activity of hiPSC-derived neuronal network grown on MEAs was recorded for 20 min. During the recording, the temperature was maintained constant at 37 °C, and the evaporation and pH changes of the medium was prevented by inflating a constant, slow flow of humidified gas (5% CO2, 20% O2, 75% N2) onto the MEA. The signal was sampled at 10 KHz, filtered with a high-pass filter (i.e. butterworth, 100 Hz cutoff frequency) and the noise threshold was set at ±4.5 standard deviations. Data analysis was performed off-line by using a custom software package named SPYCODE2 developed in MATLAB (The Mathworks, Natick, MA, USA), which allows the extraction of parameters describing the network activity.

### Data Availability

Quantified gene expression information from the RNAseq data, as described in the ‘gene expression’ heading, is available in the ArrayExpress database under accession number E-MTAB-10244. Additional intermediate data (such as tables of differential expression analysis, differentially marked genes, GO enrichments) can be found in Supplementary table 1.

## Acknowledgments

This work was funded by the Telethon Foundation (grant number GGP13231B to G.T. and GGP13231A to G.M.), the EPIGEN Flagship Project of the Italian National Research Council (to G.T.), the European Research Council (consolidator grant number 616441-DISEASEAVATARS to G.T.), the Horizon 2020 Innovative Training Network EpiSyStem (to G.T.), Ricerca Corrente granted by the Italian Ministry of Health (to G.T., G.M., and M.C.) Giovani Ricercatori granted by the Italian Ministry of Health (to G.T.), Fondazione Umberto Veronesi (to P.-L.G.), Associazione Italiana per la Ricerca sul Cancro (investigator grant to G.T. and AIRC “Fellowships for Italy” to M.G.), the Foundation IEO-CCM (fellowship to A.V.), EMBO (EMBO Short-Term Fellowship ASTF 613-2016 to M.G), the Umberto Veronesi Foundation (P.-L.G.), the Italian Ministry of Health (ERANET-Neuron grant to P.-L.G. and Ricerca Corrente grant to G.T.). This work was supported by the Netherlands Organization for Health Research and Development ZonMw grant 91217055 (to. H.v.B and N.N.K), E-Rare IMPACT consortium (G.T and H.v.B) and from the Simons Foundation (SFARI grant 610264 to N.N.K.). This research was funded by the Italian Ministry of Health, Jerome Lejeune Foundation, and Daunia Plast to G.M. S.B., S.J.K., C.M. and N.S. acknowledge the support of Newlife Charity (Grant reference 16-17/10) and Manchester University Hospitals NHS Foundation Trust, Kabuki Research Fund no. 629396. We thank the Genomic Technologies Core facility and Ian Donaldson from the Bioinformatic Core facility at the University of Manchester. We thank Kai Ge (National Institutes of Health, Bethesda, MD 20892, USA.) for sending KMT2D antibody as a gift. We are grateful to the Genomic Disorder Biobank and Telethon Network of Genetic Biobanks (Telethon Italy grant GTB12001G) and EuroBioBank (EBB) Network for banking of biospecimens and AISK (Associazione Italiana Sindrome Kabuki) for continue inspiration.

## Authors contribution

M.G. initiated this project, performed fibroblast culture, human iPSCs reprogramming, iPSCs differentiation into NCSCs, MCs. Engineered the iPSCs with *Ngn2* ePB vector and differentiation into *Ngn2* neurons, stainings for neuronal network reconstructions and MEA recordings. Performed all the immunostainings and characterization of all cell types, western blotting, and performed transcriptomic and genomic experiments in all the mentioned cell types and participated in data analysis. A.V. performed bioinformatics analysis of transcriptomic and epigenomic data of all aforementioned cell types, plus blood samples, dual SMAD samples, devised and supervised further bioinformatic analyses conducted by D. Castaldi and D. Capocefalo. S.C. produced the isogenic line and performed differentiation of dual SMAD neurons and their RNAseq library preparation. M.F.P and C.F. contributed to cell culture and cell differentiation of iPSCs, and M.F.P contributed to imaging and figure drawing. PL. G. initiated the bioinformatic analysis, and supervised A.V. during the first half of the project. D. Castaldi implemented CUT&RUN and CUT&Tag pipelines and genomics data analyses. D. Capocefalo performed re-analysis of scRNAseq of cortical brain organoids. E.T. contributed to line management and NGS library preparation. N.B.B. contributed to neuronal architecture analysis. M.F. and N.B.B. completed MEA recordings and analyzed MEA data. H.V.B., T.KL., C.S., and T.K. provided blood samples data. G.M. coordinates Telethon biobank and provided fibroblasts biopsies. N.M and G.M.S. prepared the biopsies samples for shipping to Milan. M.C and O.P. performed and analyzed SNP arrays in fibroblasts and iPSCs lines. N.N.K. supervised electrophysiological recordings and analyzed their data. S.B. and S.J.K. conceived, and along with, C.M. and N.S. designed the hESC part of the study. S.B. and S.J.K. supervised isogenic line production and dual SMAD differentiation and RNASeq data generation. G.T. supervised the whole study. M.G. and A.V. equally contributed. G.T. and G.M conceived the preliminary draft of the study, G.T., M.G., and A.V. conceived the study, interpreted data, and drafted the manuscript. All authors contributed to the final manuscript.

---

**Supplementary Figure 1.**
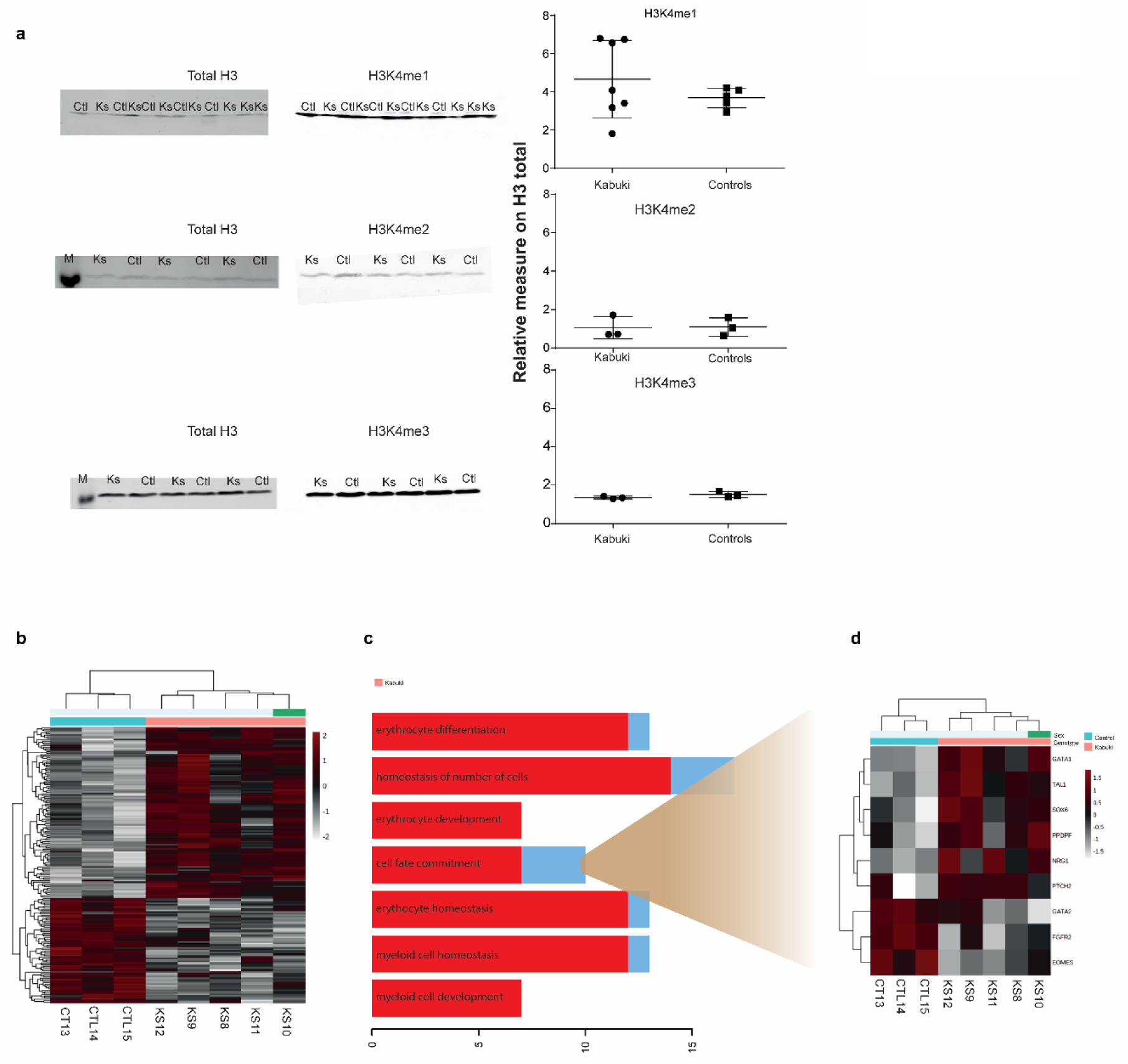
A) Western blots of bulk H3K4 mono-di- and three-methylation in KS1 fibroblasts. B) Heatmap of DEGs found in blood KS1 samples; Z-scores (log-TMM scaled by row). C) Biological Process GO enrichments of DEGs, ordered by FDR from the top; the x axis represent the number of genes; red and blue colors proportionally represent the number of up- and down-regulated genes enriching each GO term. D) Heatmap of DEGs enriching the “cell fate commitment” GO term.

**Supplementary Figure 2.**
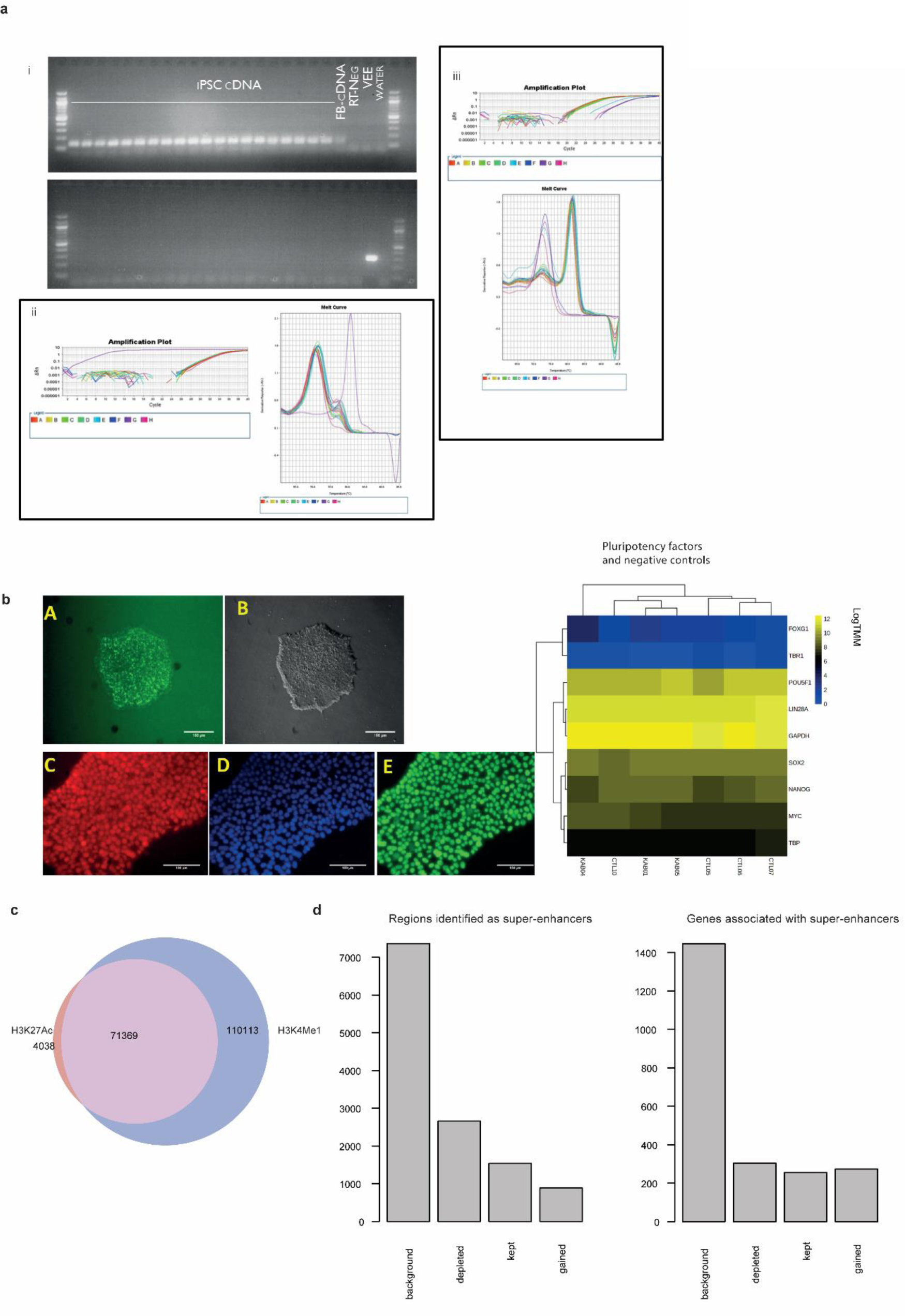
A) qPCR validation of removal for VEE: i) Upper gel: *TBP*, lower gel *VEE*; ii) Melt curve for TBP; iii) Melt curve for VEE. B) On the left: stainings of pluripotency marker in a representative iPSC colony; on the right: logTMM expression levels of pluripotency markers and negative control genes (FOXG1 and TBR1). C) Venn diagram of H3K4me1 and H3K27ac regions found in at least two samples per marker. D) Barplots of regions that show super-enhancer features in iPSCs. Background represent super-enhacers found in at least 2 samples; depleted are SE found in all CTLs and not in KS1. Kept are SE found in all CTL and all KS1. Gained are SE preferentially found in KS1. On the left: Schematics of the intersection between iPSC super-enhancer and genotype specific subsets; on the right: schematic of the intersection between genes associated to SE subsets.

**Supplementary Figure 3.**
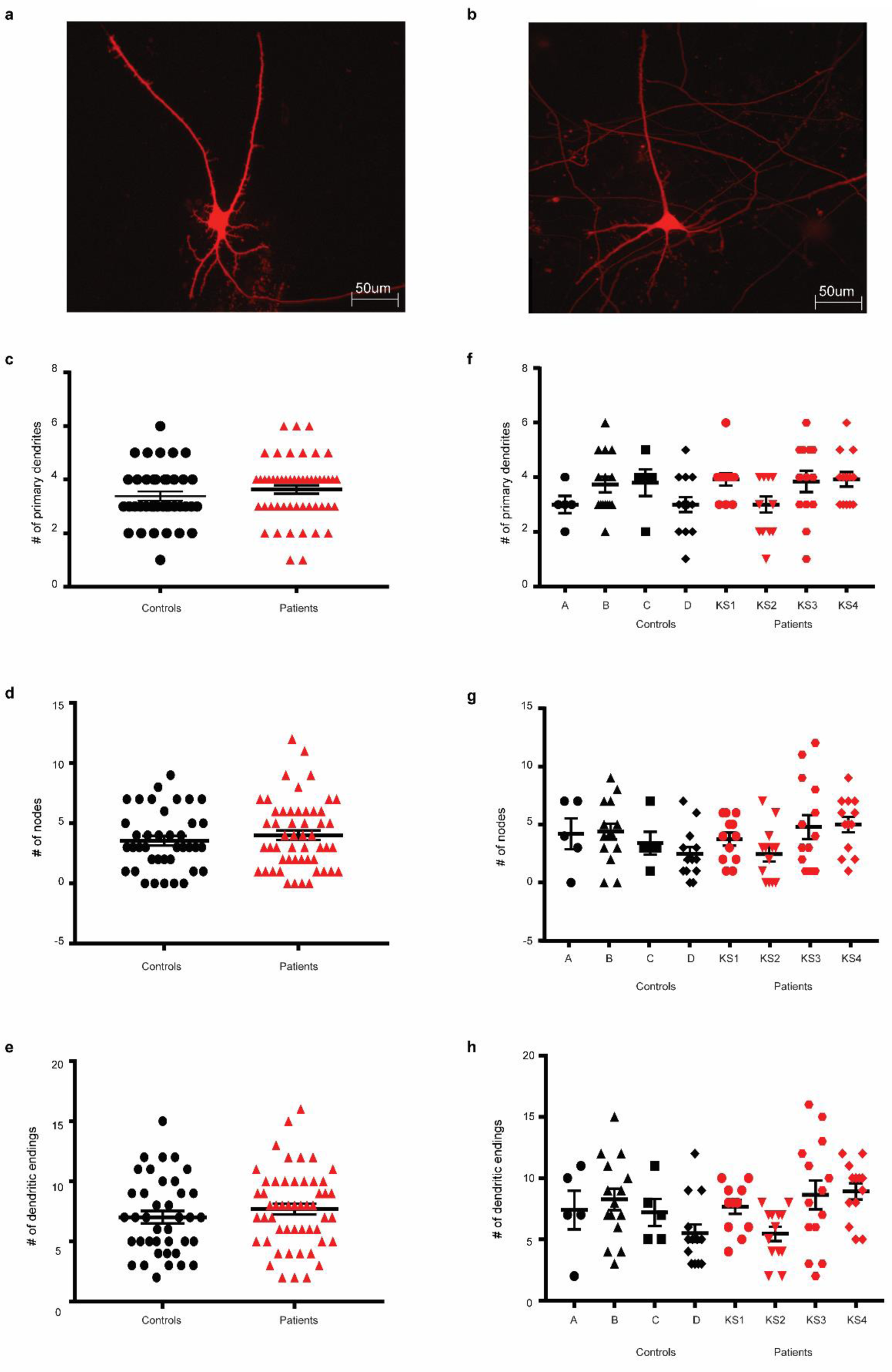
**Dendritic complexity in KS1 and control neurons.** A-B) Representative images of A) control and B) KS1 neurons transfected with dsRed (scale: 50 µm). C-E) Graphs showing the C) number of primary dendrites, D) nodes and E) dendritic endings in control (black, n=39) and KS1 neurons (red, n=52) derived from hiPSCs (i.e. pooled results). F-H) Graphs respectively showing differences in F) number of primary dendrites, G) nodes and H) endings in neurons derived from 4 control (black, C1 n=5, C2 n=15, C3 n=5, C4 n=14) and 4 KS1 (red, KS1 n=12, KS2 n=13, KS3 n=14, KS4 n=13) hiPSCs lines. Data represent means ± SEM. Statistics: normality test, Kruskal-Walis Test, post-hoc Bonferroni correction.

**Supplementary Figure 4.**
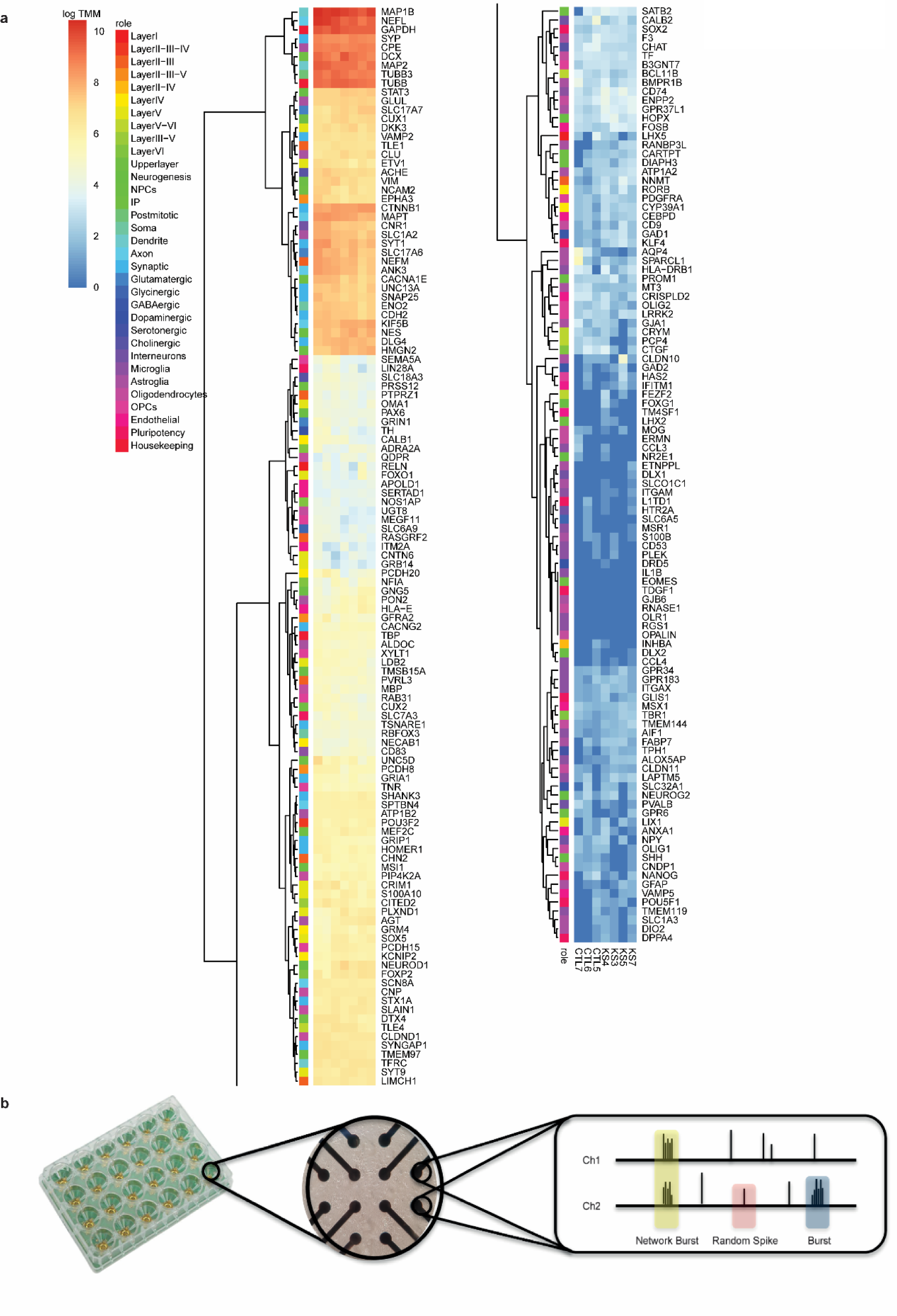
A) Expression levels (logTMM) of genes associated to different categories as markers of differentiation and neuronal development. B) Schematic of MEA logics and output.

**Supplementary Figure 5.**
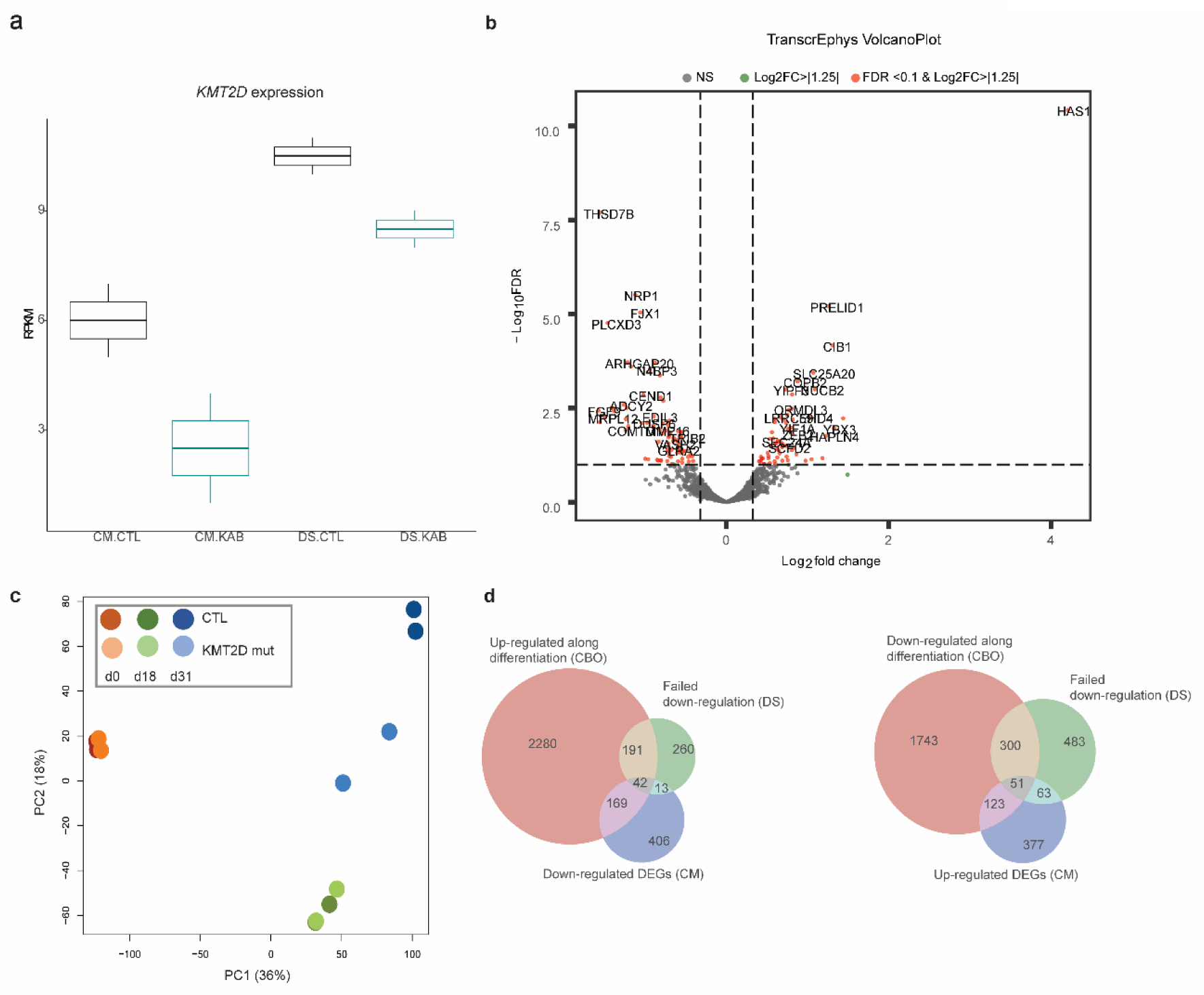
**Functional and developmental dysregulation of KS1 iNeurons** A) Boxplot of *KMT2D* levels (RPKM) in CM and DS. B) Volcano plot of transcriptional correlates of neural electrophysiology in mice, converted to human genes; log2FC and FDR measured in KS1 iNeurons. C) Principal component analysis of DS data; KS1 neurons show a day18/day31 intermediate profile; day and genotype are reported by colour as per legend; neurons (light and dark blue) show the highest inter-genotype distance. D) Venn diagrams intersecting genes whose up or downregulation failed in DS, with iN DEGs, and genes whose expression is higher (up in CBO) or lower (down in CBO) in neurons, with respect to neural precursors and radial glia, identified in single-cell RNAseq from CTL organoids.

**Supplementary Figure 6.**
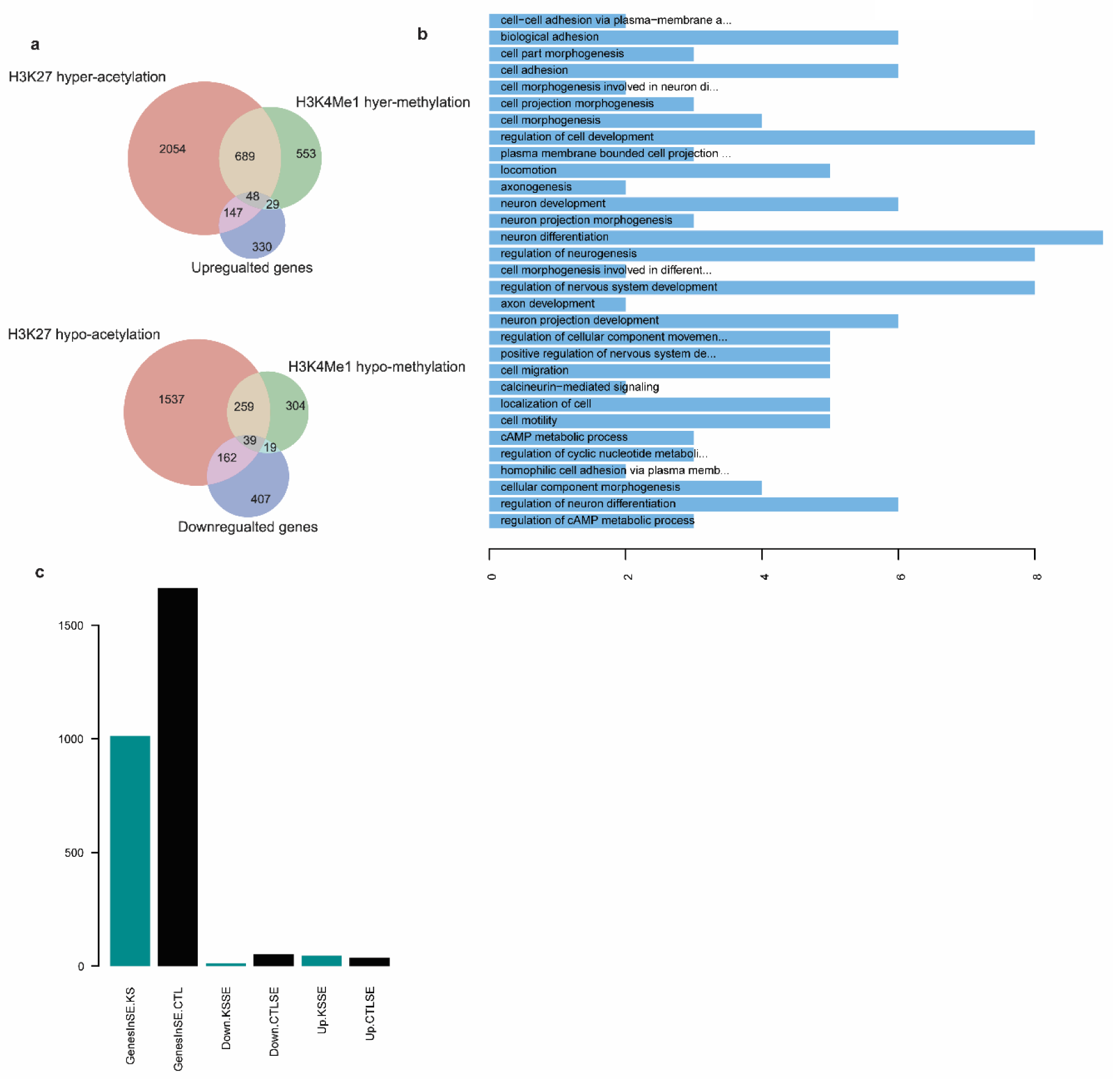
A) Venn diagrams showing the intersection between iN DEGs, and genes whose regulatory regions are differentially methylated (H3K4me1) or acetylated (H3K27ac) using p<0.05 as threshold for differentially marked regions. B) Barplots of GO enrichments for genes downregulated and hypoacetylated in KS1 iN. C) Barplots respectively depict i) genes whose regulatory regions lay within super-enhancers identified in KS1 iNeurons; ii) genes whose regulatory regions lay within super-enhancers identified in CTL iNeurons; iii) downregulated genes whose regulatory regions lay within KS1 super-enhancers; iv) downregulated genes whose regulatory regions lay within CTL super-enhancers; v) upregulated genes whose regulatory regions lay within KS1 super-enhancers; vi) upregulated genes whose regulatory regions lay within CTL super-enhancers.

**Supplementary Figure 7.**
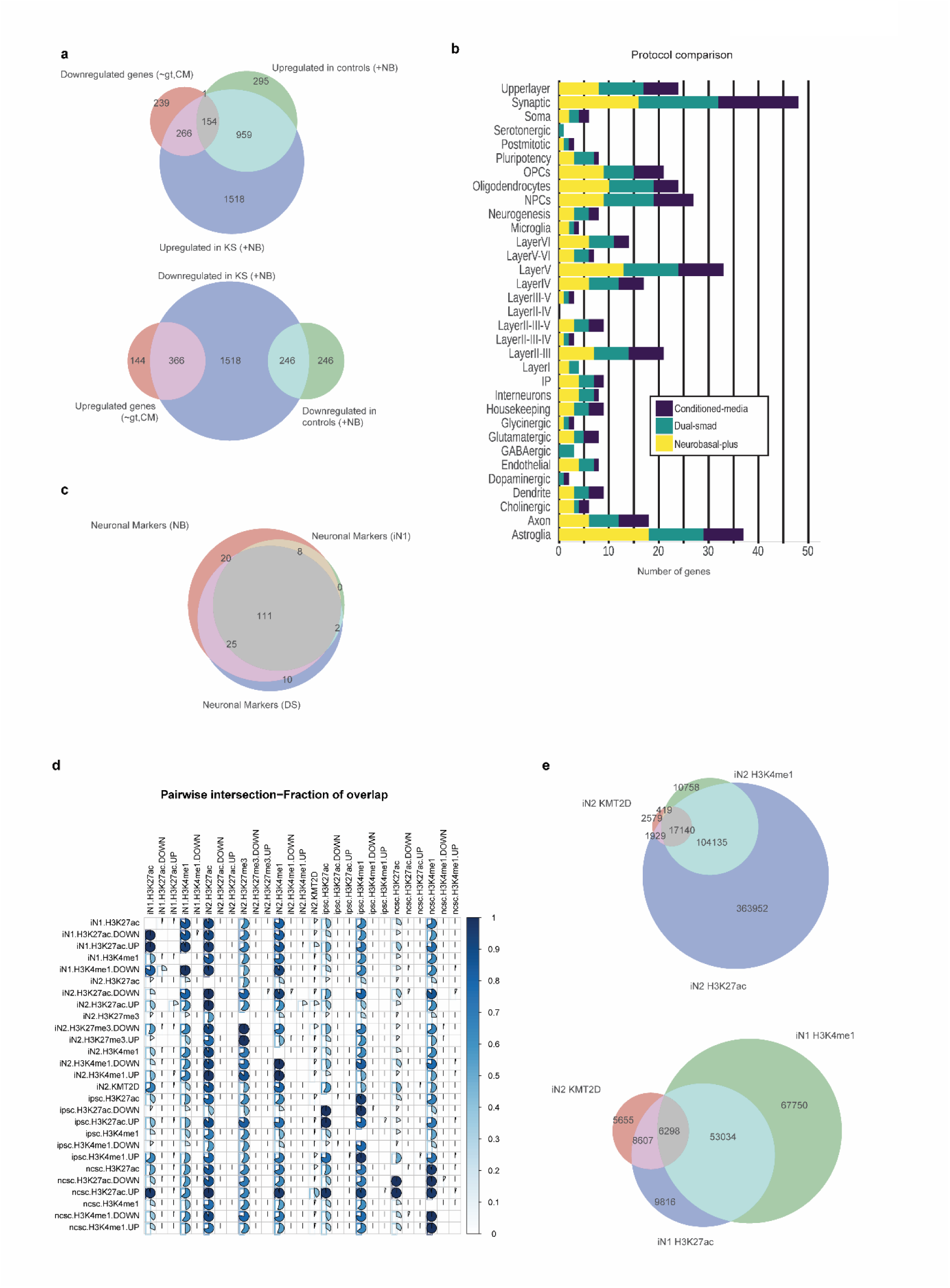
A) Venn diagram comparing genes differentially expressed in KS1 iN1 (∼gt, iN1), and genes up or downregulated in the presence of NB+ in CTL or KS1 lines. B) Stacked barplot; qualitative comparison of neuronal markers expressed in each culture condition (DS, iN1, iN2); x axis report the number of genes; each row represent a subgroup of genes. C) Venn diagram comparing neuronal markers expression in each culture condition. D) Pie heatmap of pairwise comparison between regions differentially methylated or acetylated in all tissues analyzed. E) Venn diagrams comparing genes whose enhancers are bound by KMT2D with regions marked with H3K4me1 or H3K27ac in iN2 (above) and iN1 (below).

**Supplementary Figure 8.**
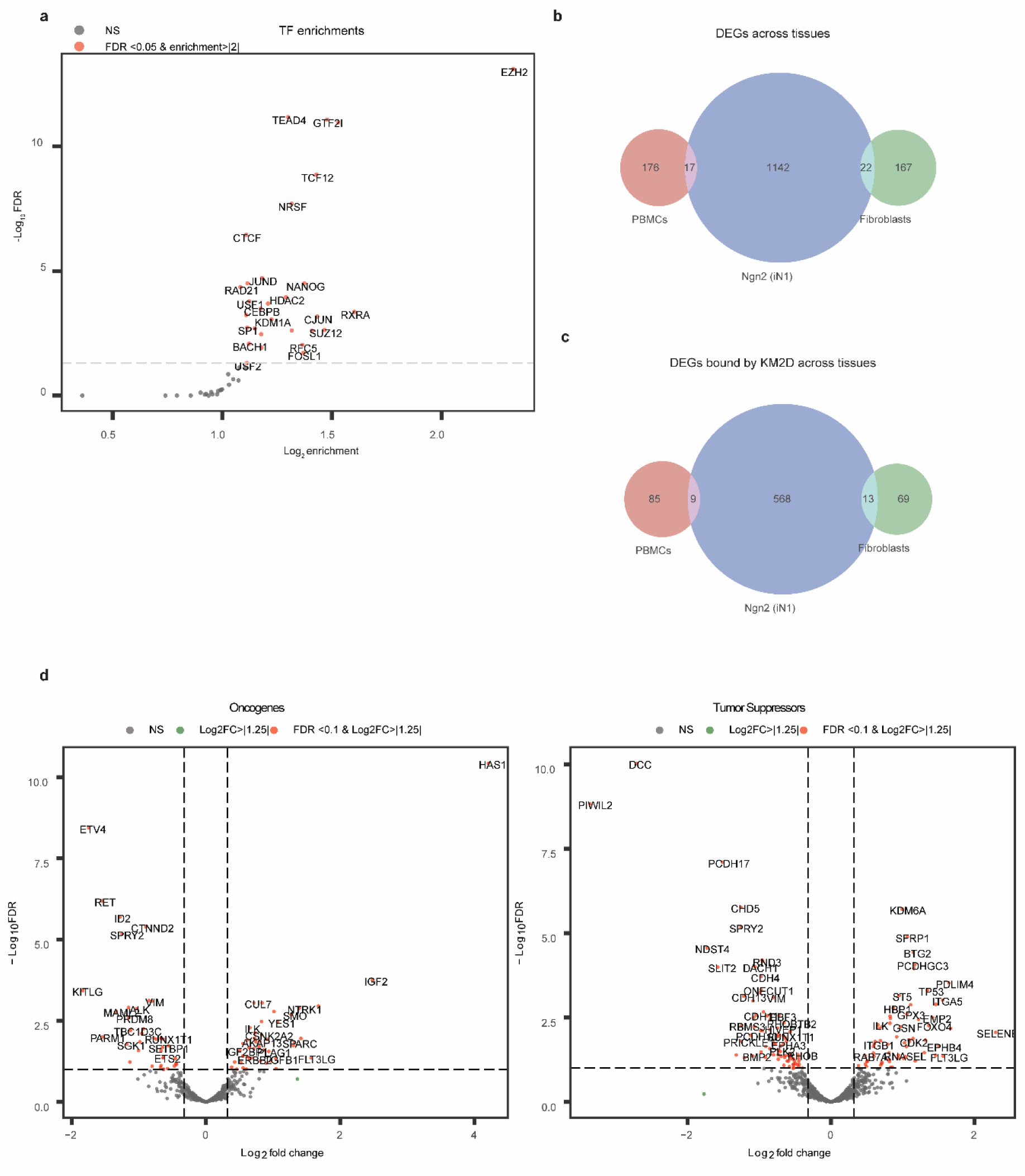
A) TF enrichment graph of differentially expressed genes in iN1 which are also bound by KMT2D at enhancers. B) Venn diagram comparing DEGs in differentiated KS1 tissues (blood, fibroblasts and iN1). C) Venn diagram comparing DEGs in differentiated KS1 tissues (blood, fibroblasts and iN1) bound by KMT2D at enhancers. D) Volcano plot of oncogenes (on the left) and tumor suppressors (on the right) differentially expressed in KS1 iNeurons.

## Footnotes

1 http://www.webgestalt.org

## Notes

### Competing Interest Statement

The authors have declared no competing interest.

